# LOSS OF PLASMA MEMBRANE LIPID ASYMMETRY CAN INDUCE ORDERED DOMAIN (RAFT) FORMATION

**DOI:** 10.1101/2021.05.03.442484

**Authors:** Shinako Kakuda, Pavana Suresh, Guangtao Li, Erwin London

**Author notes:** These authors contributed equally to this report (Shinako Kakuda, Pavana Suresh).

## Abstract

In some cases lipids in one leaflet of an asymmetric artificial lipid vesicle can suppress formation of ordered lipid domains (rafts) in the opposing leaflet. Whether suppression of domain formation might occur in plasma membranes was studied using plasma membrane vesicles (PMVs) from RBL-2H3 cells. Membrane domain formation and order was assessed by FRET and fluorescence anisotropy. Ordered domains in PMV prepared from cells by N-ethyl maleimide (NEM) treatment formed up to ~37°C, while ordered domains in symmetric vesicles formed from extracted PMV lipids were stable to 55°C, indicating that stability of ordered domains was substantially less in intact PMV. This behavior paralleled lesser ordered domain stability in artificial asymmetric lipid vesicles relative to the corresponding symmetric vesicles, suggesting that intact PMV have some degree of lipid asymmetry. This was confirmed by annexin binding showing that NEM PMV are much more asymmetric than PMV formed by dithiothreitol/paraformaldehyde treatment. Stabilization of ordered domain formation, and increased membrane order at low temperature was also observed after detergent solubilization of PMV followed by membrane reconstitution via dilution from detergent, which also should destroy asymmetry, even though membrane proteins remained associated with the reconstituted vesicles. Similar changes in domain formation and membrane order were observed after detergent reconstitution of artificial asymmetric lipid vesicles. PMV ordered domain stability was not increased by digesting peripheral domains of PMV proteins with proteinase K. We conclude loss of PMV lipid asymmetry can induce ordered domain formation. Dynamic control of asymmetry may regulate ordered domain formation in plasma membranes.

## INTRODUCTION

It is now widely accepted that the plasma membrane (PM), and likely even some internal cellular membranes, have the capacity to form co-existing liquid ordered (Lo state) lipid domains (rafts) and liquid disordered (Ld state) lipid domains (1, 2). Ordered domain formation has even been detected in bacteria, especially those with cholesterol lipids derived from host cholesterol (3, 4). Most often the formation of ordered lipid domains is driven by interactions of sphingolipids and sterol, lipids rich in PM (1). Nevertheless, much controversy remains, and how ordered domain formation, properties and size, are regulated is still unclear in most cases. Clustering of proteins with a high ordered domain affinity is one mechanism likely to promote ordered domain formation and enlargement (5–8). However, dynamic changes in lipid asymmetry also have the potential alter domain formation (9–11). This is because interleaflet coupling of lipid physical properties can alter ordered domain formation when one leaflet rich in ordered domain-forming lipids contacts a leaflet unable to spontaneously form ordered domains. Lipid asymmetry in which only one leaflet has the ability to form ordered domains can result in either induction or suppression of ordered domain formation, depending upon lipid composition (9, 10, 12–14).

Light microscopy experiments have the limitation that they cannot distinguish cases in which domain formation is suppressed from cases in which large domains are converted to submicroscopic nanodomains. Fluorescence resonance energy transfer (FRET) studies can detect nanodomains, and do not have this limitation. Using FRET, we previously demonstrated that in artificial asymmetric large unilamellar vesicles (aLUV) composed of lipids whether ordered domains are induced or suppressed is very sensitive to lipid structure (9, 10). For example, ordered domain formation is strongly suppressed in artificial asymmetric cholesterol-containing lipid vesicles mimicking PM, with an outer leaflet containing a mixture of sphingomyelin (SM) and 1-palmitoyl 2-oleoyl phosphatidylcholine (POPC), and an inner leaflet containing POPC (10). Upon loss of asymmetry in such vesicles, induction of ordered domain formation could be detected by FRET and fluorescence anisotropy (10).

The PM has a highly asymmetric lipid distribution (15), and it has been proposed that transient loss of asymmetry may have important biological functions (16). To investigate the persistence of asymmetry in isolated PM and whether there is a change in domain stability upon loss of asymmetry similar to that observed in artificial lipid vesicles, we compared domain formation in preparations of giant plasma membrane vesicles (GPMV) prepared from RBL-2H3 cells and symmetric vesicles composed of extracted PM lipids. (Note: because spectroscopic studies, unlike microscopy, report on the behavior of both the GPMV and submicroscopic vesicles present in GPMV preparations (17), we refer to the preparations used in this study as plasma membrane vesicles (PMV).)

In a previous study we demonstrated that intact PMV formed by treatment of cells with dithiothreitol (DTT) and paraformaldehyde (PFA) can form ordered domains up to physiologic temperature. In other words, they are of borderline stability at 37°C (17). We now find that PMV formed from cells by N-ethyl maleimide (NEM) treatment have a lower ability to form ordered domains than those formed by DTT/PFA treatment. However, symmetric vesicles formed from both NEM and DTT/PFA PMV lipids were found to have a strong ability to form ordered domains at and well above physiologic temperature. The difference between the behavior of NEM PMV and vesicles formed from PMV lipid was very similar to the difference between the behavior of asymmetric and symmetric artificial lipid vesicles. Annexin binding experiments showed that NEM PMV retain most of their asymmetry, and are much more asymmetric than DTT/PFA PMV. Additional experiments showed that the presence of membrane-associated proteins in the PMV cannot by itself explain the difference between intact PMV and vesicles formed from PMV lipids. We conclude that the asymmetry of PMV prepared by NEM treatment partly suppresses their ability to form ordered PM domains. Thus, there is the potential for transient loss of PM lipid asymmetry which can occur *in vivo* (11) to induce ordered membrane domain (lipid raft) formation and regulate membrane protein function.

## MATERIALS AND METHODS

### Materials

Palmitoyl-2-oleoyl-sn-glycero-3-phosphocholine (POPC), egg sphingomyelin (eSM), Cholesterol (CHOL), 1,2-dioleoyl-sn-glycero-3-phosphoethanolamine-N-(lissamine rhodamine B sulfonyl) (Rho-DOPE) were from Avanti Polar Lipids (Alabaster, AL). 1,6-diphenyl-1,3,5-hexatriene (DPH) was from Sigma-Aldrich (St. Louis, MO). Octydecylrhodamine B (ODRB), 3,3’-dilinoleyloxacarbocyanine perchlorate (FAST DiO), dithiothreitol (DTT), and 4 (w/v) % paraformaldehyde (PFA) were from Invitrogen (Carlsbad, CA). All lipids and lipid probes were stored at −20 °C. N-ethyl-maleimide (NEM) was from Sigma-Aldrich (St. Louis, MO) dissolved in methanol and stored as aliquots at −20 °C. Proteinase K (ProK) was from Thermo Fisher (Bohemia, NY). Phenylmethylsulfonyl fluoride (PMSF) was from Gold Biotechnology (St. Louis, MO). Octyl glucoside was from Anatrace (Maumee, OH) and dissolved in water. 10X phosphate-buffered saline (PBS), was from Bio-RAD (Hercules, CA). Dulbecco’s PBS (DPBS) (200 mg/l KCl, 200 mg/l KH_2_PO_4_, 8 g/l NaCl, and 2.16 g/l Na_2_HPO_4_), was from Thermo Fisher Scientific (Waltham, MA). Alexa Fluor 488 labeled Annexin V was from Life Technologies (Carlsbad, CA) and 5x annexin binding buffer from Thermo Fisher Scientific (Waltham, MA). All other chemicals were reagent grade.

### Cell culture

Rat basophilic leukemia (RBL-2H3) cells were a kind gift from Dr. Barbara Baird (Cornell University). Cells were grown in DMEM medium supplemented with 10% FBS and 100 U/ml penicillin and streptomycin and were maintained in a humidified incubator with 5% CO_2_ at 37°C.

### Preparation of giant plasma membrane vesicles (PMVs)

PMVs were produced and isolated from RBL-2H3 cells as previously described (17, 18). Briefly, cell were grown on a 10 cm plate (to 70-90 % confluent), washed twice with DPBS, and then once with PMV buffer (10 mM HEPES, 150 mM NaCl, and 2 mM CaCl_2_ (pH 7.4)). PMVs were produced by adding 2.5 ml of 2 mM NEM or 2 mM DTT/25 mM PFA in PMV buffer to the plate at 37°C for 2h. The supernatants containing PMVs were collected, and then centrifuged at 100 x g / 10 min to remove large cell debris. The resulting PMV-containing supernatants were used for the measurements on intact PMV (e.g. in Figure 1 and 2) within two hours of preparation. Because we found the PMV induced by NEM results in contamination with small cell debris or cytosolic components that resulted in high backgrounds when measuring DPH fluorescence (see below), in Figure 1 and 2 the background intensities were subtracted for the FRET measurements. For subsequent experiments, PMVs were pelleted by further centrifugation at 16,100 x g at 4°C for 30 min to obtain higher concentrations needed in later lipid extraction and reconstitution experiments. After this step, the background averaged around 2% of the DPH fluorescence intensity and was not subtracted from DPH fluorescence. Analysis of the crude PMVs by flow cytometry (see below) showed the preparations were contaminated by at most 1-2% of large cell debris (Supplemental Figure 1). There was more debris in NEM preparations than in the DTT/PFA preparations. In both cases, the debris was reduced several fold by the centrifugation at 100 x g. It should also be noted that contamination by residual cell debris would not be expected to increase ordered domain formation. Overall, cell lipids have a lower SM content than PM, and a lower tendency to form an ordered bilayer (Supplemental Figure 2). Thus, contamination of PM preparations by other cell membranes would imply the tendency of PM lipids to form ordered domains is even higher than we report here.

**Figure 1.**
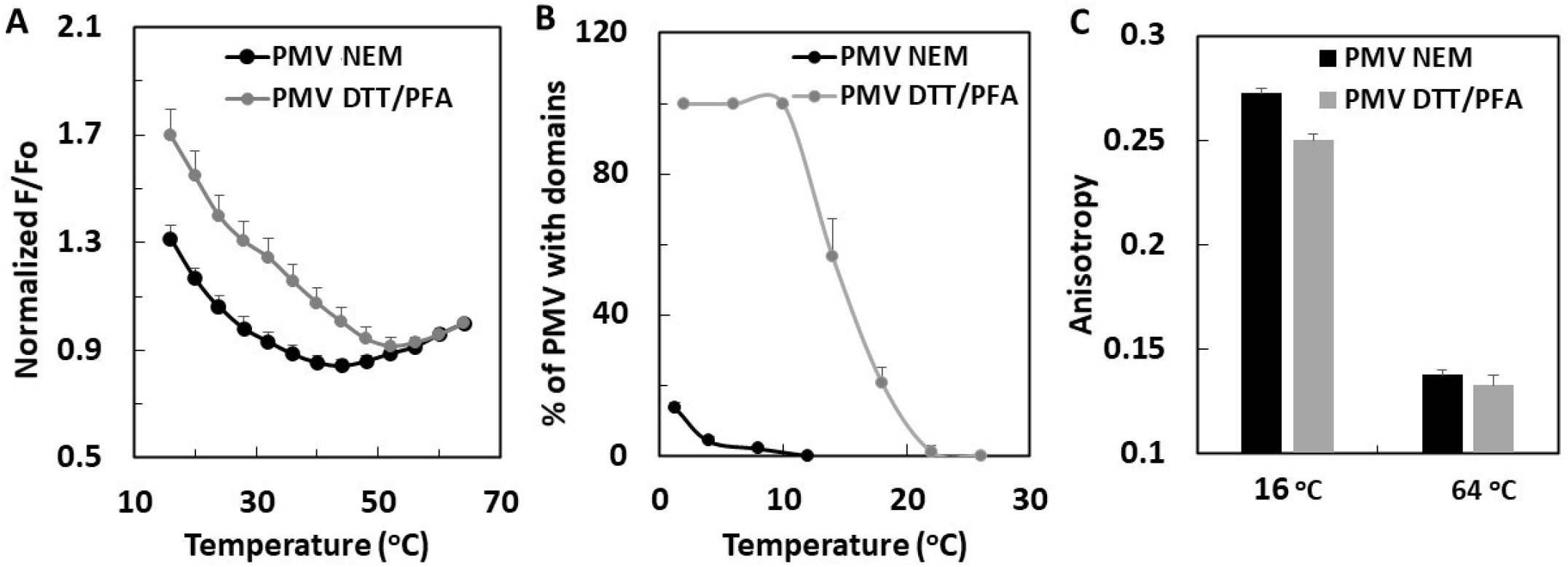
Temperature dependence of ordered domain formation in PMV from RBL-2H3 cells. (A) Nanodomain formation detected by FRET. DPH (FRET donor) and ODRB (FRET acceptor) were added to F samples. Only DPH added to Fo samples. F/Fo values normalized to 1 at 64 °C. (B) Phase separation detected by fluorescence microscopy. Cells labeled with FAST DiO before induction of PMV. (C) Fluorescence anisotropy of membrane-inserted DPH. Gray lines and bars are PMV prepared by DTT-PFA, black lines and bars are PMV prepared by NEM. Unless otherwise noted, in this and following figures mean values and standard deviations (only up direction bar is shown) from three independent preparations are shown. For this and following figures unnormalized F/Fo vs. temperature FRET curves are shown in Supplemental Figures.

**Figure 2.**
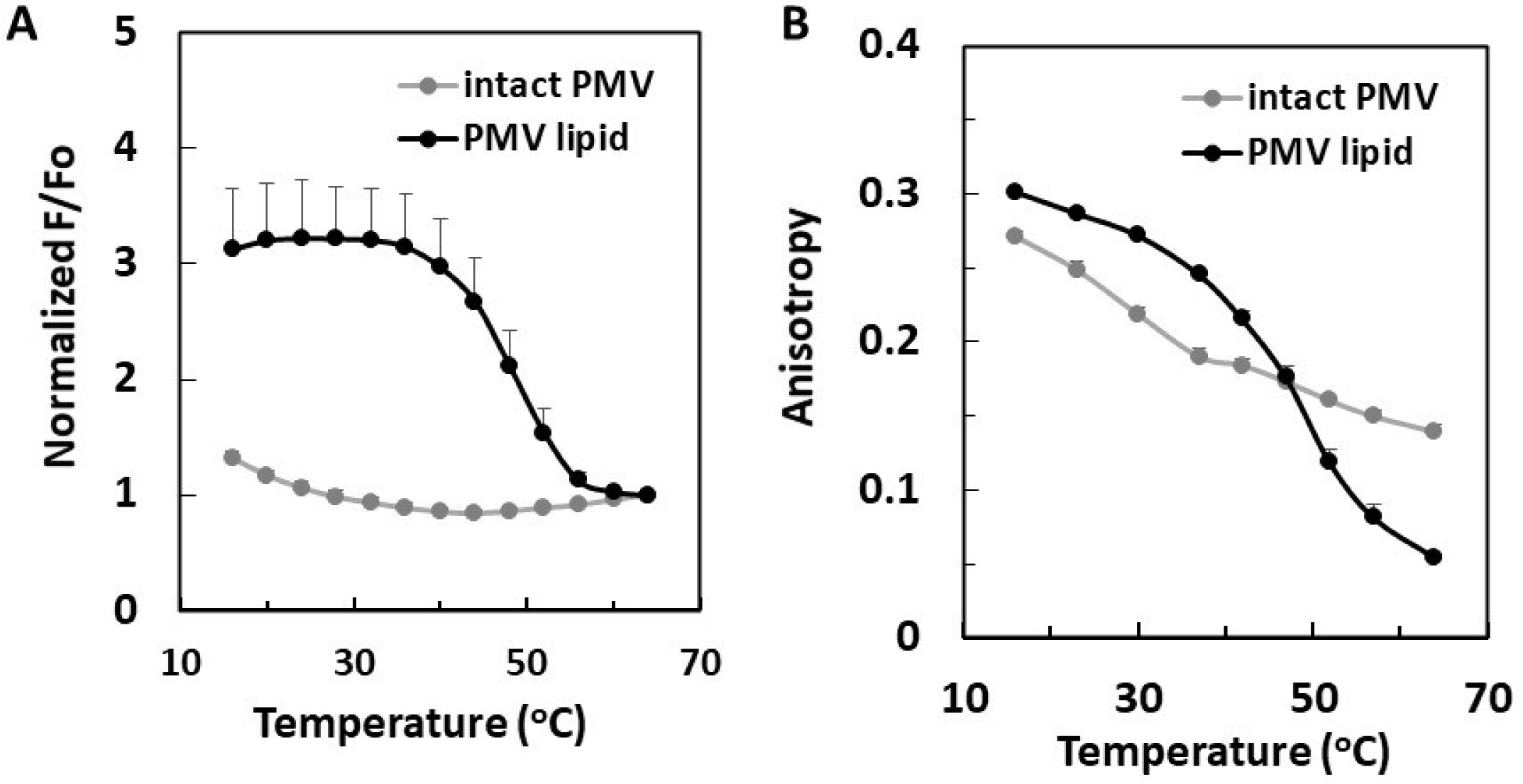
Comparison of domain formation and membrane order in intact PMV and lipid vesicles prepared from extracted PMV lipids. In this and subsequent figures PMV prepared using the NEM method. (A) Temperature dependence of nanodomain formation detected by FRET. (B) Temperature dependence of membrane order by fluorescence anisotropy of membrane-inserted DPH: (gray line) intact PMV, (black line) lipid vesicles extracted from intact PMV.

### Annexin Staining of PMV and Flow Cytometry

PMVs were produced as described above with 2 mM NEM or 5 mM DTT/25 mM PFA from a 10 cm plate. (5 mM DTT was used for these experiments as this maximized the difference between annexin binding by NEM and DTT/PFA PMV). For these experiments, the PMVs were collected after the centrifugation at 16,100xg at 4°C for 30 min. One third of this PMV pellet was resuspended in 100 μl of annexin binding buffer (ABB, 10 mM HEPES, 140 mM NaCl, 2.5 mM CaCl2) and 5 μl of Annexin V Alexa Fluor 488 (Life technologies corporation, CA) was added and samples incubated for 15 min in the dark. Then, 400 μl of ABB was added and the samples filtered using a 35 μm mesh cap (Falcon) and then used for flow cytometry. To prepare a crude PMV pellet, PMVs from the 10 cm plate were pelleted at 16,100 x g at 4°C for 30 min and the pellet was resuspended in 500 ul ABB and filtered before using for flow cytometry. Flow cytometry was performed using a Cytoflex LX (Beckman Coulter, CA), collecting approximately 50,000 events per sample. Data was analyzed using FlowJo version 10 software (FlowJo LLC).

### Microscopy of Annexin Stained PMV

PMVs from RBL-2H3 cells stained with annexin V AF-488 as described above were allowed to settle for 1h in the chamber of MatTek Glass Bottom Microwell Dishes pre-coated with 1% BSA (P35G-1.5-10-C, MatTek corporation). The PMV were visualized using ET-EGFP (FITC/Cy2) filters (Chroma) and AF-488 fluorescence recorded using Olympus IX83 fluorescence microscope. Raw fluorescence intensity values were used for analysis of PMV fluorescence intensity and processed using ImageJ. To do that, for each PMV fluorescence intensity along a line crossing the equator was analyzed with the Plot Profile program. Intensity at the pixel with highest fluorescence minus background intensity was then recorded. For each condition, the average fluorescence intensity was calculated from 25 PMVs of similar size. The ratio of intensity in DTT/PFA PMV to that in NEM PMV was then calculated.

### Phase separation detected by fluorescence microscopy

Large-scale domain formation was visualized by staining cells with Fast DiO as previously described (17). Briefly, cells were stained with 5 μg/ml Fast DiO/ethanol at 4°C for 10 min and then produced PMV as above. The resulting Fast DiO labeled PMVs were pelleted by centrifugation at 500 x g for 10 min and then suspended in 100-500 μl of PMV buffer for microscopy imaging. The fraction of PMV having domains were assessed in 150 PMVs from three images as previously described (17).

#### Preparation of PMV lipids

PMV pellets (stored at −80°C for up to a few days) from two to three 10 cm plates were extracted by 0.8-1.2 ml of 3 : 2 (v : v) hexanes : isopropanol at room temperature for 30 min. Under these conditions lipid extraction was complete as shown by a lack of lipid that could be detected by thin layer chromatography after a second extraction step. Thin layer chromatography was carried out as previously described (19).

### Preparation of lipid vesicles from extracted PMV lipids

After extraction of lipid, the organic solvent was dried under a nitrogen stream and either used that day, or after overnight storage at −20°C. The lipids were hydrated with 2 ml of PMV buffer for FRET measurements or about 0.2 ml of PMV buffer for octyl glucoside reconstitution, and proteinase K digestion experiments (see below). The lipid mixtures were dispersed at 70°C for 5 min and then vortexed at 55°C for 30 min using a multi-tube vortexer. The resulting multi-lamellar vesicles (MLVs) were subjected to seven cycles of freezing-thawing using liquid nitrogen-warm water to preparing symmetric vesicles from the PMV lipids. These vesicles were used the same day they were prepared or after storage overnight at 4°C.

### Preparation of artificial symmetric large unilamellar vesicles (sLUV) and asymmetric large unilamellar vesicles (aLUV)

Artificial symmetric and asymmetric lipid vesicles were prepared as previously described (9, 10). For comparisons, symmetric vesicles were prepared similarly to asymmetric vesicles, with entrapped sucrose, and pelleted prior to use. The mol % composition of sLUV was 22.5 : 52.5 : 25 eSM : POPC : cholesterol; and of aLUV was about 45 : 30 : 25 eSM : POPC : cholesterol outer leaflet and 75 : 25 POPC : cholesterol inner leaflet (10).

### Measurement of PMV lipid concentration in vesicles by DPH

Lipid concentration of PMVs and artificial lipid vesicles were estimated by measuring the fluorescence intensity of membrane-inserted DPH under conditions in which DPH is in excess (17, 20). A standard curve of bSM : POPC 1:1 SUV was prepared. To do this, 0.5 μmol bSM and 0.5 μmol POPC were mixed and dried under a nitrogen stream. The dried lipids were dissolved in 30 μl ethanol and then 970 μl of PBS was added, and incubated at 70°C for 5 min to disperse the lipids. Next 5 μl of 0.2 mM DPH/ethanol was added to the desired dilution of the standard in PBS (1 ml total volume) to give a range of 0-15 μM of SUV. (It should be noted that because we were interested in the relative lipid concentration in different PMV preparations, no correction for any lipid composition dependence of DPH fluorescence was made.) Fluorescence was measured in the experimental samples (generally diluted to 0.5-3 μM lipid) after incubation for 5 min at room temperature in the dark, The intensity of DPH fluorescence in standard curve and experimental samples were measured using λex 358 nm; λem 427 nm and slit bandpass of 3 nm.

### Measurement of the temperature dependence of FRET

FRET measurements on PMV and vesicles made from PMV lipids were performed with DPH as the FRET donor and ODRB as the FRET acceptor. Samples contained 10-16 μM lipid dispersed in 1 ml of PMV buffer. For “F samples” 4 μl of 100 μM ODRB/ethanol was then added. “Fo samples” lacked ODRB. Both types of samples were then incubated at 37°C for 1 h in the dark. Next, 4 μl of 7.5 μM DPH/ethanol was added to both F samples and Fo samples, and incubated at room temperature for 5 min in the dark. DPH fluorescence intensity was measured using a Horiba QuantaMaster Spectrofluorimeter (Horiba Scientific, Edison, NJ) using quartz semimicro cuvettes (excitation path length 10 mm and emission path length 4 mm. DPH fluorescence was measured at λex 358 nm; λem 427 nm, and bandpass set 3 nm for both excitation and emission). When desired, background fluorescence intensity (i.e. in the absence of DPH) was measured from readings for both F samples and Fo samples. Data was collected at every 4°C while increasing temperature from 16°C to 64°C. Between readings temperature was increased at a rate of 4°C/5 min. The fraction of fluorescence not quenched by FRET is equal to F/Fo. (FRET efficiency is 1-F/Fo). When necessary (data in Figures 1 and 2), background values were subtracted before F/Fo was calculated. In later experiments in which the PMV were pelleted by centrifugation to remove contaminants, the background was negligible (average about 2% of signal). F/Fo ratios were normalized to 1 at 64°C to compensate for variations arising from variable incorporation of ODRB into PMV. Unnormalized F/Fo data is shown in the Supplemental Materials. It should be noted the background arising from contaminants was higher in PMV prepared with NEM than those prepared with DTT/PFA, probably due to easy detachment of cells during PMV formation when using NEM.

For artificial vesicles composed of SM-POPC-CHOL, the FRET measurement protocol was similar except instead of ODRB, samples contained as FRET acceptor rhodamine-DOPE (3% of total lipid) which was mixed with the lipids in organic solvent before drying. Again, final lipid concentration was 10-16 μM dispersed in 1 ml of PBS. FRET measurements were carried out the same day the vesicles were prepared.

### Octyl Glucoside (OG) treatment of PMV and artificial vesicles

Pellets of PMV (freshly made), lipid vesicles from PMV lipids, or artificial lipid vesicles, were suspended in a small volume of PBS, to give a concentration of about 300-450 μM lipid. Then OG/water was added to give a final concentration 45 mM OG. The samples were then incubated at 37°C for 1h, followed by dilution to 1.5 mM OG and 10 – 15 μM lipid to reconstitute vesicles. For control, 1.5 mM OG was added to PMV and lipid vesicle samples containing 10 – 15 μM lipid.

### Fluorescence anisotropy measurements

Fluorescence anisotropy was measured as previously described (10). Lipid concentrations were 10-15 μM. DPH was 0.03 μM, which gives a DPH/lipid ratio of 1/300 to 1/500.

### Proteinase K (ProK) treatment

A 200 μl aliquot of PMV with 100-160 μM lipid was incubated with or without 8 μg of ProK in PMV buffer at 37°C for 1 h with gentle vortexing every 15 min. ProK activity was then inhibited by adding 2 mM phenylmethylsulfonyl fluoride (PMSF) from a stock solution in ethanol, for 15 min at room temperature. The resulting ProK treated PMVs were diluted to 2 ml for FRET measurement as above. For checking ProK efficiency, 20 μl of untreated and of ProK treated PMV (before dilution for FRET) was loaded to 8 % SDS-PAGE gels and the proteins were stained with GelCold Blue Stain (Thermo Fisher Scientific). A control in which PMSF was added for 15 min at room temperature before adding ProK also prepared and loaded on the gel.

## RESULTS

### Ordered domain formation in NEM-induced and DTT/PFA-induced PMV

Control of ordered domain formation in PMV was studied. FRET and fluorescence anisotropy were used to assess physical properties sensitive to ordered lipid domain formation and lipid asymmetry (9, 10, 21). Relative to microscopy, FRET has the advantage that it can detect nanodomains as small as 5-10 nm. In the FRET assay, the fraction of fluorescence of the FRET donor (DPH) unquenched by the presence of a rhodamine-containing FRET acceptor (octadecylrhodamine B (ODRB) or rhodamine-DOPE (9, 17, 22) was measured (Ro about 3.6 nm (22)). This fraction is equal to F/Fo, where F is DPH fluorescence in the presence of acceptor and Fo is DPH fluorescence in the absence of acceptor. (FRET efficiency is 1-F/Fo.) The FRET acceptors used preferentially partition into disordered domains (7), while the FRET donor DPH generally partitions nearly equally between ordered and disordered domains (18). The consequence is that FRET is weaker (F/Fo higher) in membranes containing co-existing ordered Lo domains and disordered Ld domains than it is in homogenous membranes lacking Lo domains (9, 17, 22). At high temperatures, at which ordered domains “melt”/become miscible with Ld lipids, there is a decrease in the average distance between DPH molecules that were localized in the ordered domains and FRET acceptors that were in the disordered domains. As a result, overall F/Fo decreases. The thermal stability of ordered domains can be estimated by the temperature at the apparent midpoint (inflection point) in the roughly sigmoidal F/Fo versus temperature curves. In cases in which the transition is at too low a temperature, thermal stability can be estimated by the approximate endpoint temperature, at which the absolute value of the slope of F/Fo vs. temperature reaches a minimum (17, 23).

Fluorescence anisotropy measurements were carried out to confirm FRET results. The fluorescence anisotropy of membrane-inserted DPH, a measure of membrane order, is higher in the Lo state (~0.3) than the Ld state (~0.05-0.15)(24).

The temperature dependence of domain formation in PMV prepared from cultured RBL-2H3 cells by two different methods was compared. The first method was treatment of cells with a combination of dithiothreitol (DTT) and paraformaldehyde (PFA), and the second treatment with N-ethyl maleimide (NEM)(25). The DTT/PFA method is widely used because the large-scale domains in DTT/PFA-induced PMV can be easily visualized at up to almost room temperature by light microscopy, whereas large scale domains in NEM-induced PMV can be detected only at very low temperature (25). However, DTT/PFA is a much more perturbing chemical treatment than NEM, as it crosslinks proteins and chemically modifies phosphatidylethanolamine (PE), a major PM lipid.

Comparison of nanodomain formation in NEM- and DTT/PFA-induced PMV vs. temperature, as assayed by FRET, is shown in Figure 1A. (Unnormalized FRET data is shown in Supplemental Figures 2,4-6). The nanodomains in NEM-induced PMV were significantly less thermally stable than nanodomains in DTT/PFA-induced PMV. This difference paralleled that for large scale domain formation, which disappeared above 5-10 °C in NEM-induced PMV and 20 °C in DTT/PFA-induced PMV (Figure 1B), roughly in agreement with previous studies (25). As noted previously, DTT/PFA-induced PMV nanodomain formation is more thermally stable than large scale phase separation (17). Figure 1 shows this is also true for NEM-induced PMV.

The anisotropy of NEM- and DTT/PFA-induced PMV at 16 °C and 64 °C was also compared (Figure 1C). Both preparations exhibited high anisotropy at low temperature, consistent with a high level of ordered domain formation, and low anisotropy indicative of Ld state at high temperature, indicating loss of ordered domain formation at high temperature. The difference between ordered domain stability in the two preparations does not seem to reflect a different amount of SM relative to PC in the NEM-induced PMV relative to that in the DTT/PFA-induced PMV (Supplemental Figure 3). Instead, a difference in asymmetry might be responsible for the difference in ordered domain thermal stability in the NEM-induced and DTT/PFA-induced PMV (see below). Unless otherwise noted, in subsequent experiments, the less chemically-perturbed NEM-induced PMV were studied.

### Ordered domain formation in PMV is enhanced in vesicles prepared from PMV lipid extracts

We recently reported that loss of lipid asymmetry induced ordered lipid domain formation in artificial vesicles that roughly mimic PM (10). Therefore, we hypothesized that if NEM-PMV retained a significant level of asymmetry, symmetric lipid vesicles composed of NEM-PMV lipids would form more stable ordered domains than those in intact PMV. To test this hypothesis, we prepared artificial large unilamellar vesicles (which exhibit lipid symmetry (26–28)) from extracted NEM-PMV lipids, and compared ordered domain stability in these vesicles to that in intact PMV. (It should be noted that although PMV lipid vesicles were prepared by freeze-thaw without subsequent extrusion, they were largely unilamellar as judged by lipid accessibility to external reagents, see Supplemental Figure 7.)

As predicted, the FRET curve demonstrated that, ordered domains formed in PMV lipid were considerably more thermally stable, persisting to 55°C, than those in intact PMV, which disappeared above about 37°C (Figure 2A). (Ordered domain formation in lipid extract vesicles from PMV made with DTT/PFA treatment was similar to that for vesicles prepared from PMV made with NEM treatment, see Supplemental Figure 8). FRET results were confirmed by DPH fluorescence anisotropy (Figure 2B). An anisotropy vs. temperature curve closely paralleled the FRET curve for PMV lipid, with the highly ordered state being lost above 55°C. Note that a weaker temperature dependent decrease in anisotropy continues above the temperature at which the domains melt. This decrease occurs because a disordered state bilayer becomes increasingly disordered as temperature is increased. Anisotropy in intact PMV also showed a domain melting event in which ordered domains disappeared (at about 37°C), similar to that detected by FRET. It is noteworthy that at low temperature DPH anisotropy in intact PMV was lower than that for PMV lipid vesicles (Figure 2B). This suggests a lower extent of ordered domain formation in intact PMV than in PMV lipid at low temperature. Overall, the conclusions from FRET and anisotropy are in good agreement.

### Lipid asymmetry of PMV preparations

The results described above suggest the possibility that, as is the case in artificial vesicles with a simple lipid composition (9, 10), the loss of asymmetry present in NEM PMV can induce/enhance ordered domain formation. To assess NEM PMV asymmetry binding of annexin-V, which binds to phosphatidylserine (PS), a lipid that is located on the inner leaflet in the plasma membrane of intact cells, was measured (29). It should be noted that prior studies found only a small fraction (~10%) of PMV are permeable to molecules the size of annexin-V (16) Prior studies have also shown that PMV formed by DTT/PFA treatment lose at least some of their asymmetry (30). Therefore, we compared annexin binding by NEM PMV to annexin binding by DTT/PFA PMV. As shown in Figure 3A, microscopy detected more (about five-fold higher) annexin-V binding to DTT/PFA than to NEM PMV. Similar results were obtained using flow cytometry (Figure 3B and 3C) including when comparing the entire PMV preparation, or excluding the small fraction of debris, or comparing PMV fractions of equal size as judged by forward scattering levels (Supplemental Figure 9). Assuming the DTT/PFA PMV are fully symmetric, and that permeability to annexin V can be ignored this would suggest that NEM PMV retain close to 80% of their PS asymmetry. If DTT/PFA PMV retain some asymmetry, this could be an underestimate of NEM PMV asymmetry.

**Figure 3:**
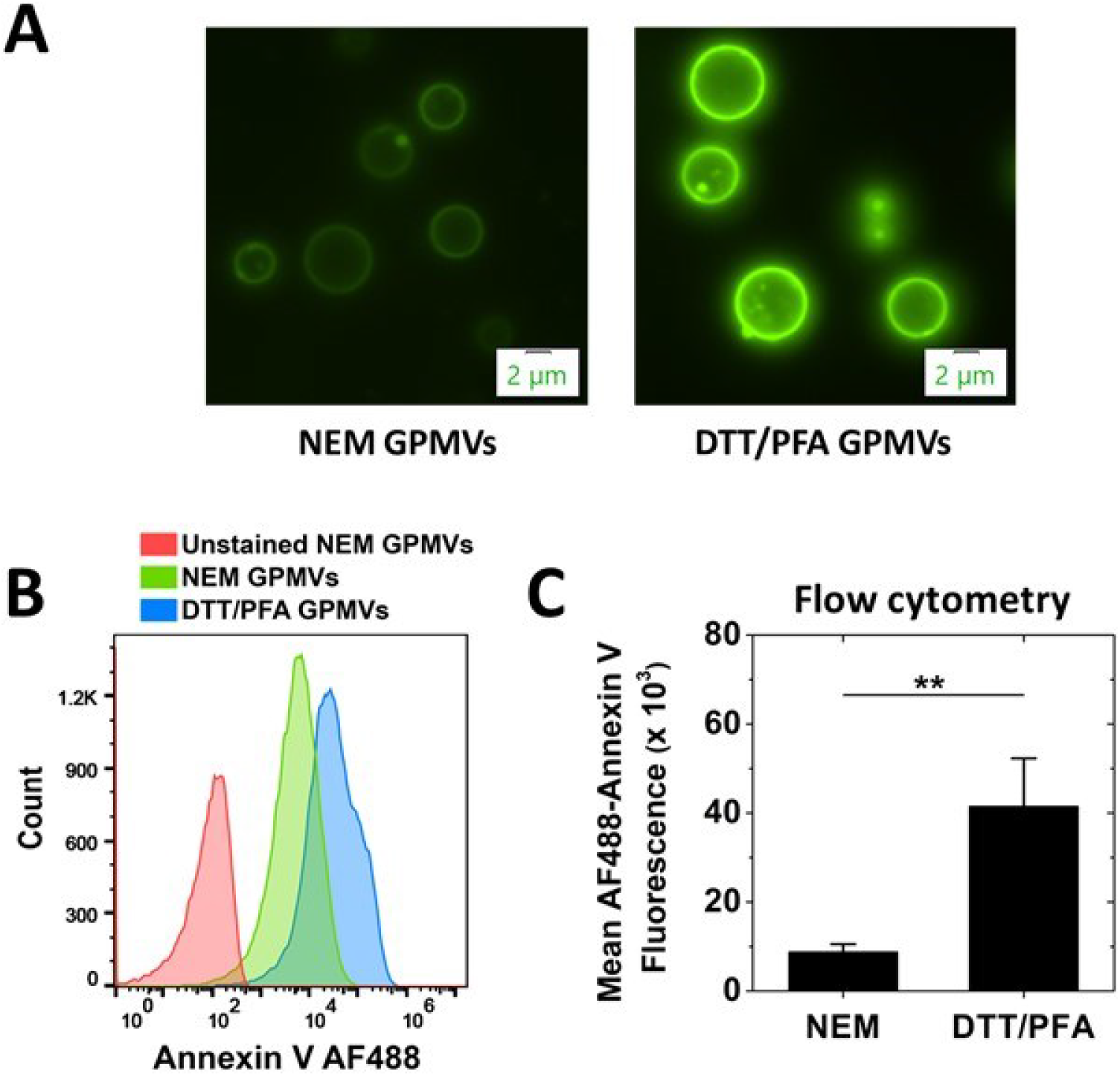
RBL-2H3 PMVs made using 2 mM NEM and 5 mM DTT/25 mM PFA stained with Annexin V-AF488. (A) Micrographs of PMVs stained with Annexin V-AF488 at the same microscope exposure settings. The Annexin V stain intensity for DTT/PFA PMVs was 5.1-fold higher than for NEM PMVs. (B) Flow cytometry of gated PMVs showing number of PMV counts vs Annexin V-AF488 binding to PMVs for unstained sample (red), NEM PMVs (green), and DTT/PFA PMVs (blue). Fluorescence from about 50,000 PMVs is shown. Notice x-axis is logarithmic. (C) Mean fluorescence intensity of NEM and DTT/PFA PMVs from flow cytometry data is shown from three experiments with about 50,000 events counted per experiment. **p<0.01. The ratio of intensity of DTT/PFA PMVs to NEM from flow cytometry experiments was 4.96 ± 1.95.

### The role of asymmetry and proteins in ordered domain formation in PMV

Another possibility to explain the difference between intact PMV and vesicles prepared from PMV lipid is that proteins associated with intact PMV are responsible for the difference in behavior of intact PMV and that of vesicles formed from extracted PMV lipids. To examine this possibility, and further test the effect of loss of lipid asymmetry, we examined whether treating intact PMV under conditions that should destroy asymmetry, but not remove membrane proteins, would destabilize ordered domain formation. To do this vesicles were dissolved in detergent and then the detergent was diluted to a concentration well below its critical micelle concentration to re-form vesicles (31, 32).

To confirm that detergent treatment destroyed asymmetry, we first studied the effect of detergent solubilization and dilution using octyl glucoside (OG) upon asymmetric artificial lipid vesicles that crudely mimic PM. The behavior of asymmetric large unilamellar vesicles (aLUV) with SM and POPC in the outer leaflet, POPC in the inner leaflet and cholesterol in both leaflets, was compared using FRET to that of ordinary symmetric unilamellar vesicles (sLUV) composed of SM-POPC-cholesterol. As shown in Figure 4A, after OG solubilization of aLUV and reconstitution (to form symmetric vesicles), ordered domain formation was increased/thermally stabilized relative to untreated aLUV samples, This difference in behavior is very similar to what we reported previously for sLUV and aLUV for this lipid composition (10). Control experiments simply adding OG to aLUV at the final OG concentration (OG control) showed little effect on FRET. Figure 4B shows that there was also little effect on ordered domain formation and stability when the symmetric SM-POPC-cholesterol vesicles (with about the same overall composition as the aLUV) were first dissolved in OG and then vesicles reconstituted by dilution, as expected as vesicles should remain symmetric after reconstitution. Therefore, stabilization of ordered domain formation upon solubilization and reconstitution is specific to the asymmetric vesicles. This is consistent with our previous report showing that a loss of asymmetry triggered by heating vesicles can induce ordered domain formation in aLUV composed of SM, POPC and cholesterol (10).

**Figure 4.**
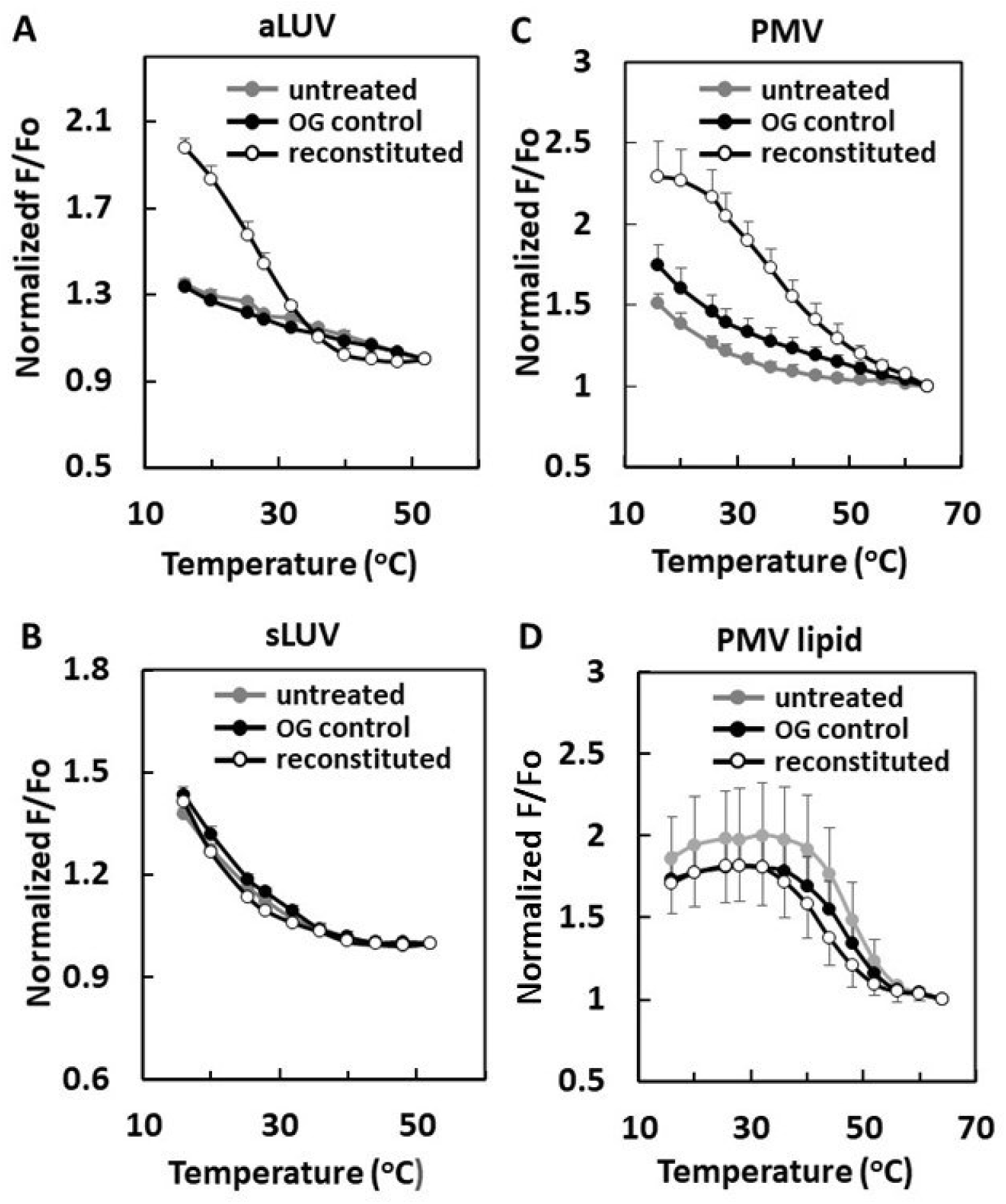
Effect of OG reconstituted membranes upon domain formation. Samples contained: (A) aLUV of SM-POPC outside/POPC inside/cholesterol (see the composition ratio in methods); (B) sLUV of SM-POPC-cholesterol (see the composition in methods); (C) intact PMV; (D) PMV lipids. FRET in vesicles PMV derived samples used DPH as donor and ODRB as acceptor. FRET in artificial LUV used DPH as donor and rhodamine-DOPE as acceptor. FRET measured on: (gray circles) untreated vesicles; (open circles) reconstituted vesicles prepared by dissolving vesicles in 45 mM OG and then diluting to 1.5 mM OG; (black circles) vesicles to which 1.5 mM OG was added as a control (OG control). For clarity, in 3D. standard deviations for reconstituted vesicles are illustrated as down bars.

Analogous experiments were then carried out upon intact PMV prepared with NEM treatment and extracted PMV lipid vesicles (Figures 4C and 4D). FRET showed that reconstitution of PMV after solubilization with OG increased ordered domain stability to a level similar to that in the vesicles composed of PMV lipids. This was not due to a loss of protein upon solubilization and reconstitution. This was shown by sucrose gradient centrifugation of OG reconstituted vesicles formed from intact PMV compared to that for OG reconstituted vesicles formed from PMV lipids. As shown in Supplemental Figure 10, the vesicles formed from PMV had a higher density than the vesicles from just PMV lipids, and protein co-migrated with lipid in the PMV-derived samples. This implies the change in ordered domain stability was not due to loss of protein. The fact that domain stability was not altered in controls in which OG was added to vesicles at the final concentration (Figure 4C) indicates OG itself was not responsible for the increase in ordered domain formation. The observation that there was almost no change in ordered domain stability after reconstituting OG-solubilized vesicles composed of PMV lipid (Figure 4D), which should already be symmetric before OG reconstitution, indicates that the change in domain formation after OG reconstitution of intact PMV reflects a change in asymmetry. This conclusion is further supported by the observation that domain formation after OG reconstitution of PMV lipid vesicles, was similar to that of OG reconstitution of intact PMV (Figure 4C).

DPH fluorescence anisotropy was measured to strengthen the conclusion that upon reconstitution of OG-solubilized artificial aLUV there was increased ordered domain stability associated with a loss of asymmetry. Figure 5A shows that untreated SM-POPC-cholesterol sLUV containing ordered domains exhibited a high degree of anisotropy at low temperature (16°C), and OG reconstitution of sLUV had no significant effect on membrane order (Figure 5A, left). In contrast, untreated aLUV with about the same overall lipid composition as the sLUV had a much lower level of anisotropy than untreated sLUV (Figure 5A, right), consistent with the FRET result indicating very little ordered domain formation in aLUV. However, after solubilization and reconstitution, the anisotropy levels of aLUV increased to that found in sLUV (Figure 5A, left). Adding OG directly to aLUV (or sLUV) or at the final concentration present after reconstitution induced at most a small decrease of anisotropy, rather than an increase (Figure 5C). These experiments confirm the FRET results, showing that OG solubilization and dilution to destroy asymmetry in aLUV induces domain formation.

**Figure 5.**
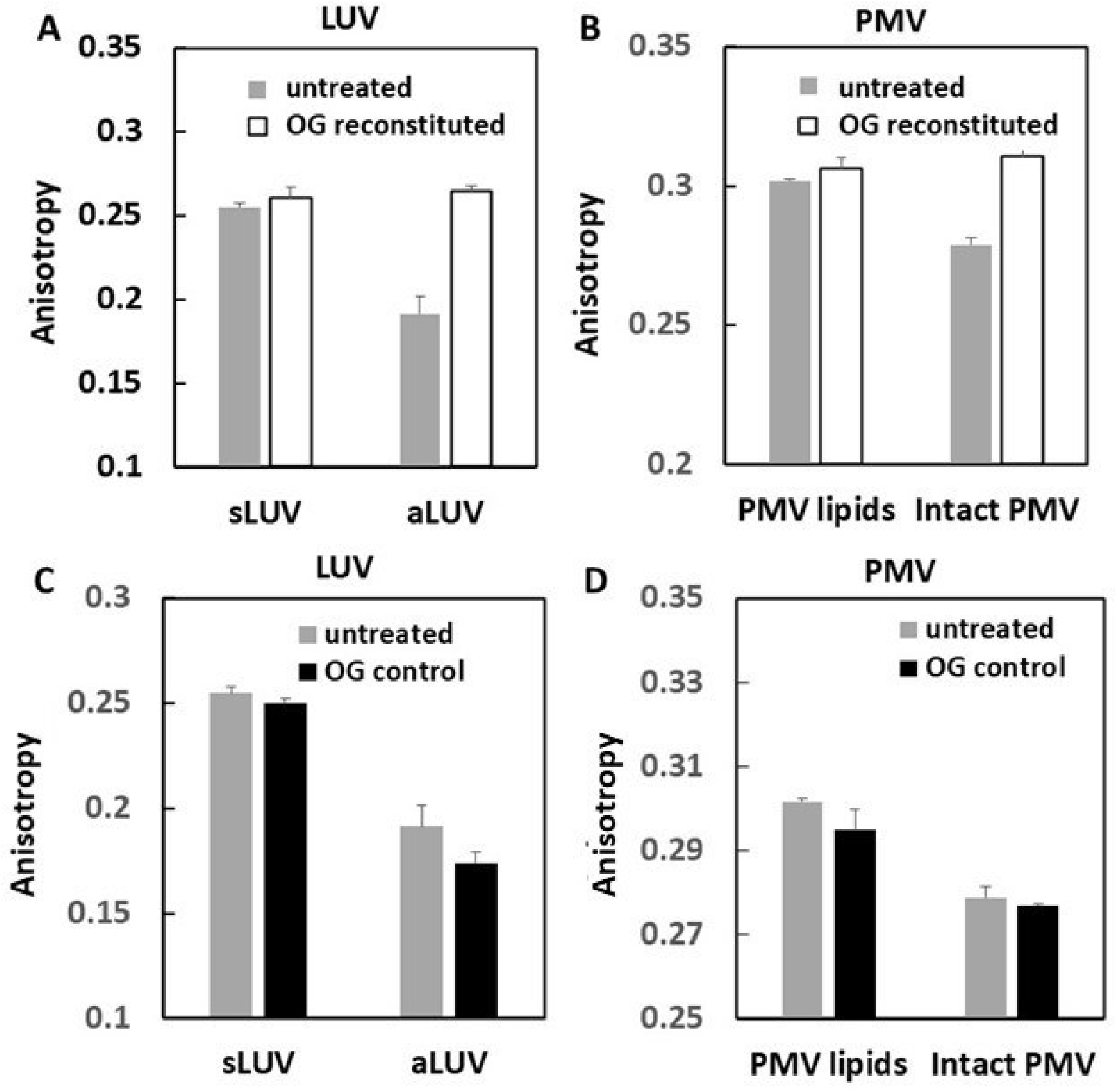
Reconstitution of intact PMV increases membrane order. (A and C) Artificial LUV, (B and D) PMV derived vesicles. Anisotropy of DPH fluorescence was measured at 16°C on: (gray bars) untreated vesicles; (open bars) reconstituted vesicles prepared by dissolving lipid vesicles in 45 mM OG and then diluting to 1.5 mM OG; (black bars) vesicles to which 1.5 mM OG was added as a control (OG control).

Figure 5B shows analogous experiments for vesicles formed from PMV lipid and intact PMV. Vesicles from PMV lipid, and those from PMV lipid re-formed after OG solubilization and reconstitution of those vesicles, both exhibited a similar high degree of anisotropy (Figure 5B, left), as expected for vesicles that are symmetric. In contrast, intact PMV had a lower level of anisotropy than that of the vesicles formed after solubilization and reconstitution of PMV (Figure 5B right), as expected if asymmetric before solubilization and symmetric after solubilization. This was true even though the vesicles reconstituted after solubilization of intact PMV contain PMV proteins, as noted above. Again, direct addition of OG to PMV (or PMV lipid) vesicles at the final concentration after reconstitution had little effect on or slightly decreased anisotropy (Figure 5D, right). Thus, the effect of vesicle solubilization and reconstitution upon DPH fluorescence anisotropy in intact PMV and vesicles formed from PMV lipid agrees with the FRET results for PMV and vesicles from PMV lipids in terms of showing a loss of asymmetry increases membrane order, as well as with the anisotropy changes seen after reconstitution of artificial asymmetric and symmetric lipid vesicles, which show a loss of asymmetry increases membrane order in artificial vesicles.

Table 1 schematically summarizes the properties of the artificial vesicles and PMV. Notice that the differences in both domain forming properties (assayed by FRET) and membrane order properties (as measured by anisotropy) of PMV and PMV lipid vesicles, and how they are affected by reconstitution, closely parallel the differences between these properties in asymmetric and symmetric artificial vesicles, and how they are affected by reconstitution.

**TABLE 1:**
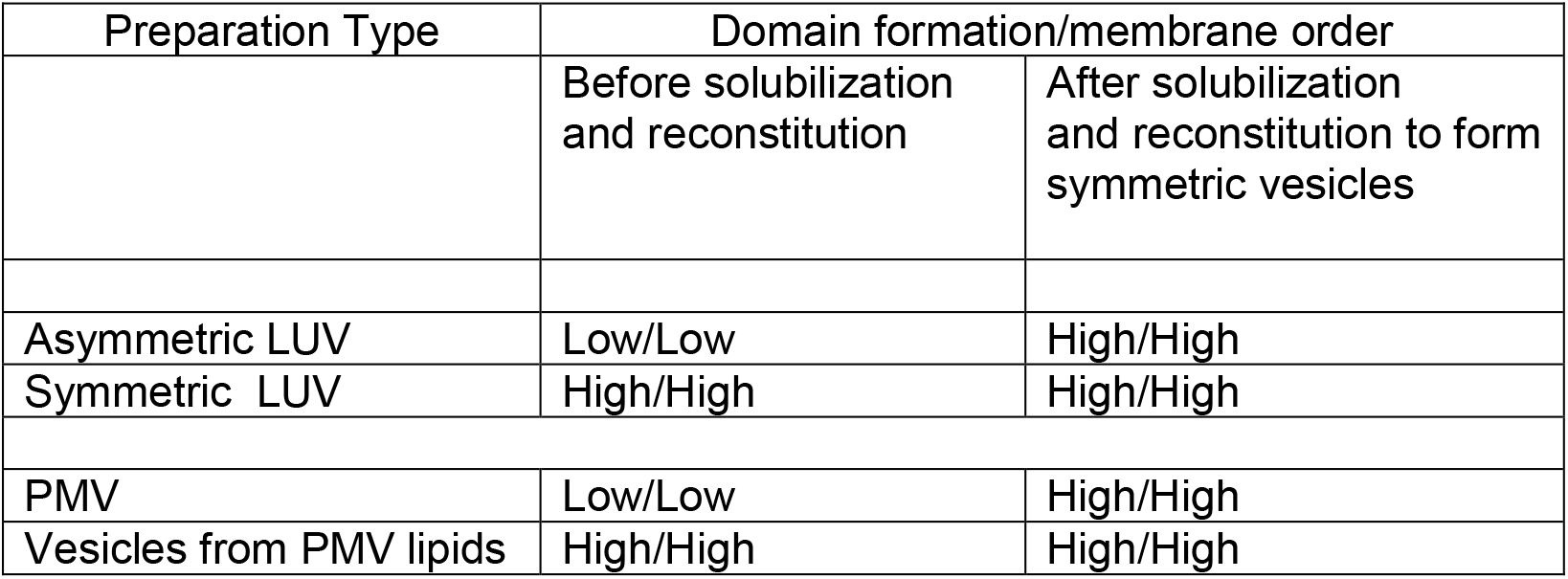
Schematic Summary of Comparison of Properties of Artificial Lipid Vesicles to that of PMV/PMV Lipid Vesicles.

To further investigate if PMV proteins suppress ordered domain formation in intact PMV, PMVs were treated with proteinase K (ProK), a non-specific protease that strongly degrades at least the non-transmembrane domains of proteins (33). There was no increase in the thermal stability of ordered domain formation as assayed by FRET (Figure 6A) and no large change in DPH fluorescence anisotropy after digestion with ProK (Figure 6B) despite extensive degradation of proteins (Figure 6C). If there was any significant change, ProK treatment appeared to reduce, not to increase, membrane order slightly, at least at elevated temperature.

**Figure 6.**
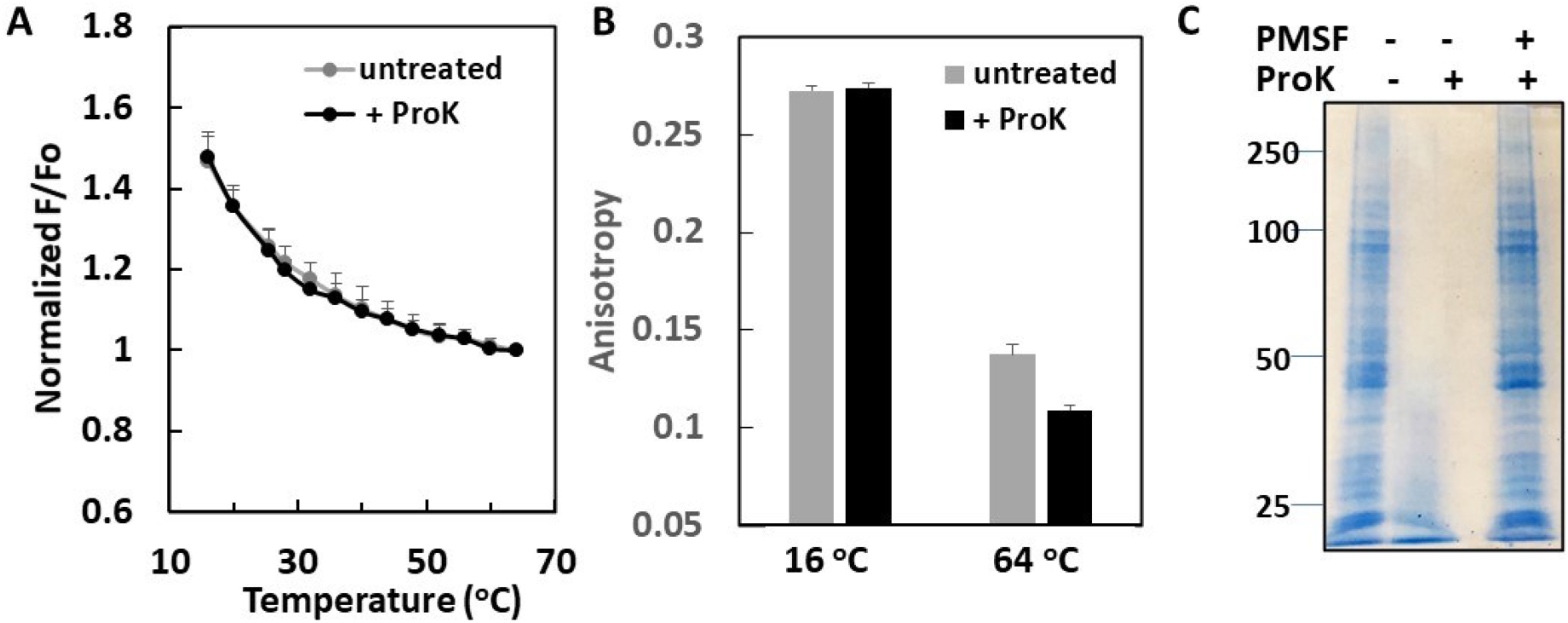
Effect of ProK treatment of intact PMV upon domain formation. (A) FRET measurements of (gray line) untreated and (black line) ProK treated intact PMV. FRET donor was DPH and acceptor was ODRB. (B) Anisotropy of (gray bars) untreated and (back bars) ProK treated intact PMV at 16 °C and 64 °C. (C) Example of SDS-PAGE analysis of the efficiency of ProK treatment. (left lane) untreated, (middle lane) ProK treated, (right lane) control for ProK treatment

Thus, both FRET and anisotropy data strongly support the conclusion that ordered domain formation in intact PMV is suppressed by lipid asymmetry, and that loss of this asymmetry stabilizes ordered domain formation. However, this does not prove proteins have no effect on ordered domain formation (see Discussion).

## DISCUSSION

### Capacity of PMV and PM lipids to form ordered membrane domains and dependence upon lipid asymmetry and proteins

The studies in this report show that plasma membrane lipids have an ability to form ordered lipid domains (rafts) that is suppressed in the natural plasma membranes. Several lines of evidence showed lipid asymmetry was one factor that suppresses domain formation. We measured physical properties that assess domain formation and membrane order, which several studies have shown are also sensitive to lipid asymmetry (9, 10, 21, 34). The studies show that PM lipids in symmetric lipid vesicles have a strong ability to form co-existing ordered and disordered lipid domains. The ordered domains were stable to well above physiologic temperature. In contrast, intact NEM PMV formed ordered domains with much lower thermal stability. Based on prior studies that show asymmetry can repress ordered domain formation (9, 10), this suggested that intact NEM PMV have considerable lipid asymmetry, as was confirmed by much lower annexin V binding to PS than for DTT/PFA PMV. A role for asymmetry was further supported by the observations that:

1. detergent reconstitution of NEM PMV to destroy asymmetry increased membrane order at low temperature and ordered domain stability to levels seen in symmetric PM lipid vesicles, and 2. NEM PMV and symmetric vesicles formed from PMV lipids had differences in properties very similar to the differences between asymmetric and symmetric SM-POPC-cholesterol lipid vesicles. This was true for differences between: a. domain formation, b. membrane order and c. response to reconstitution. The observation that DTT/PFA-induced PMV have lost much more asymmetry than NEM-induced PMV and that ordered domains formed by DTT/PFA-induced PMV are more thermally stable than those by NEM-induced PMV also is consistent with this hypothesis.

Our studies did not detect large effects of proteins on ordered domain formation. Loss of cytoskeletal connections is associated with PMV formation (30), and membrane domains are more easily detected in PMV than intact cells. Therefore, it is possible that residual cytoskeletal connections in PMV inhibit ordered domain formation relative to that in vesicles prepared from PMV lipid. However, the observation that proteinase K digestion did not alter domain formation indicates this is not what inhibited PMV ordered domain formation in the PMV. Another possibility is that transmembrane segments alter domain formation. Overall, TM segments in PM proteins have amino acid sequences in contact with the outer leaflet lipids that favor contact with Lo state lipid, and sequences in contact with the inner leaflet lipids that favor interaction with Ld state lipid (36). In addition, proteinase K does not digest all TM segments (33). However, there was no destabilization of domain formation in the protein-containing vesicles reconstituted from intact PMV by detergent dilution. This indicates that protein-induced suppression of ordered domain formation does not explain our observations. Nevertheless, in cells proteins can certainly influence domain formation; as noted above clustering of proteins with a high ordered domain affinity is one mechanism likely to promote ordered domain formation and enlargement (5–8). We would not rule out the possibility that membrane proteins make some contribution to the suppression of ordered domain formation in PMV.

It should be noted that we used different FRET acceptors for artificial lipid vesicles and PMV or PMV lipid vesicles. In previous studies of artificial lipid vesicles (9, 10, 23) we used rhodamine-DOPE, which can be mixed with other lipids in organic solvent before vesicles are prepared, and continued to use that probe in artificial lipid vesicles in this study to allow comparison to previous studies. However, rhodamine-DOPE is too water insoluble to add to preformed vesicles, so we use the rhodamine-containing probe ODRB in PMV, and to allow comparison, also used it in lipid vesicles formed from PMV lipids. Since we are not directly comparing the artificial lipid vesicles to ones containing PMV lipids, this should not be of concern. In addition, comparison of FRET using ODRB and rhodamine-DOPE in artificial vesicles, shows that they detect similar ordered domain stabilities (Supplemental Figure 11).

### Potential Implications of changes in asymmetry for ordered domain formation *in vivo*

The observation that loss of PM asymmetry can induce formation of stable ordered domains under physiologic conditions at which they would not otherwise form, or would be of only borderline stability, is of potential biological significance because transient loss of asymmetry in PM is associated with a number of biological events, including signal transduction (11). *In vivo,* it is likely that activation of lipid scramblases, such as TMEM16F, are responsible for the loss of lipid asymmetry (35). TMEM16F is activated by cytosolic Ca^2+^, and cytosolic Ca^2+^ concentration becomes elevated during signal transduction (35). This could induce ordered domain formation. It is also possible that ordered domain formation could be induced in PM prior to loss of asymmetry, upon clustering of proteins that favor formation of Lo domains, or upon changes in PM interactions with cytoskeletal proteins. In that case, the loss of lipid asymmetry might be part of a positive feedback loop, in which receptor activation and clustering induces or enhances ordered domain formation (5–8), followed by increases in cytosolic Ca^2+^ that activate lipid scramblases, resulting in a loss of asymmetry that further enhances ordered domain formation.

### Implications for interpretation of detergent resistant membrane formation from cells

It is known that lipids in the Lo state can be detergent insoluble, and that the detergent resistant membranes (DRM) isolated from cells (37) are at least largely in the Lo state (36, 38, 39). However, DRM are not likely to be identical to the ordered domains present in cells in the absence of detergent. The most widely used protocols involve incubation with Triton X-100 at 4°C, and low temperature could induce the formation of ordered domains not present at physiological temperature (1). In addition, based on studies in artificial symmetric lipid vesicles, it has been proposed that the Triton X-100 bound to membranes can induce lateral segregation of Lo and Ld domains (40), although our prior studies indicate instead that Triton X-100 only induces coalescence of preexisting ordered nanodomains (22). In any case, the present studies suggest that a loss of asymmetry could be additional issue that must be considered. Because detergent can induce lipid flip-flop, which would destroy asymmetry (41, 42), a detergent-induced loss of lipid asymmetry might enhance DRM formation upon detergent treatment of cells.

## Data Availability Statement

Data generated or analyzed during this study are included in this published article (and its supplementary information files) or are available from the corresponding author on reasonable request.

## Acknowledgement

This work was supported by NIH grant GM 122493 to E.L.

## References

1. Brown, D.A., and E. London. 2000. Structure and function of sphingolipid- and cholesterol-rich membrane rafts. The Journal of biological chemistry 275: 17221–17224.

2. Wang, H. Y., D. Bharti, and I. Levental. 2020. Membrane Heterogeneity Beyond the Plasma Membrane. Front Cell Dev Biol 8: 580814.

3. LaRocca, T. J., J. T. Crowley, B. J. Cusack, P. Pathak, J. Benach, E. London, J. C. Garcia-Monco, and J. L. Benach. 2010. Cholesterol lipids of Borrelia burgdorferi form lipid rafts and are required for the bactericidal activity of a complement-independent antibody. Cell Host Microbe 8: 331–342.

4. LaRocca, T. J., P. Pathak, S. Chiantia, A. Toledo, J. R. Silvius, J. L. Benach, and E. London. 2013. Proving lipid rafts exist: membrane domains in the prokaryote Borrelia burgdorferi have the same properties as eukaryotic lipid rafts. PLoS Pathog 9: e1003353.

5. Day, C. A., and A. K. Kenworthy. 2015. Functions of cholera toxin B-subunit as a raft cross-linker. Essays Biochem 57: 135–145.

6. Sezgin, E., I. Levental, S. Mayor, and C. Eggeling. 2017. The mystery of membrane organization: composition, regulation and roles of lipid rafts. Nat Rev Mol Cell Biol 18: 361–374.

7. Buenaventura, T., S. Bitsi, W. E. Laughlin, T. Burgoyne, Z. Lyu, A. I. Oqua, H. Norman, E. R. McGlone, A. S. Klymchenko, I. R. Correa, Jr., A. Walker, A. Inoue, A. Hanyaloglu, J. Grimes, Z. Koszegi, D. Calebiro, G. A. Rutter, S. R. Bloom, B. Jones, and A. Tomas. 2019. Agonist-induced membrane nanodomain clustering drives GLP-1 receptor responses in pancreatic beta cells. PLoS Biol 17: e3000097.

8. Chung, J. K., W. Y. C. Huang, C. B. Carbone, L. M. Nocka, A. N. Parikh, R. D. Vale, and J. T. Groves. 2021. Coupled membrane lipid miscibility and phosphotyrosine-driven protein condensation phase transitions. Biophysical journal 120: 1257–1265.

9. Wang, Q., and E. London. 2018. Lipid Structure and Composition Control Consequences of Interleaflet Coupling in Asymmetric Vesicles. Biophys J 115: 664–678.

10. St Clair, J. W., S. Kakuda, and E. London. 2020. Induction of Ordered Lipid Raft Domain Formation by Loss of Lipid Asymmetry. Biophys J 119: 483–492.

11. Doktorova, M., J. L. Symons, and I. Levental. 2020. Structural and functional consequences of reversible lipid asymmetry in living membranes. Nat Chem Biol 16: 1321–1330.

12. Lin, Q., and E. London. 2015. Ordered raft domains induced by outer leaflet sphingomyelin in cholesterol-rich asymmetric vesicles. Biophys J 108: 2212–2222.

13. Collins, M. D., and S. L. Keller. 2008. Tuning lipid mixtures to induce or suppress domain formation across leaflets of unsupported asymmetric bilayers. Proceedings of the National Academy of Sciences 105: 124–128.

14. Wan, C., V. Kiessling, and L. K. Tamm. 2008. Coupling of cholesterol-rich lipid phases in asymmetric bilayers. Biochemistry 47: 2190–2198.

15. Verkleij, A., R. Zwaal, B. Roelofsen, P. Comfurius, D. Kastelijn, and L. Van Deenen. 1973. The asymmetric distribution of phospholipids in the human red cell membrane. A combined study using phospholipases and freeze-etch electron microscopy. Biochimica et Biophysica Acta (BBA)-Biomembranes 323: 178–193.

16. Skinkle, A. D., K. R. Levental, and I. Levental. 2020. Cell-Derived Plasma Membrane Vesicles Are Permeable to Hydrophilic Macromolecules. Biophys J 118: 1292–1300.

17. Li, G., Q. Wang, S. Kakuda, and E. London. 2020. Nanodomains can persist at physiologic temperature in plasma membrane vesicles and be modulated by altering cell lipids. Journal of lipid research 61: 758–766.

18. Sezgin, E., H. J. Kaiser, T. Baumgart, P. Schwille, K. Simons, and I. Levental. 2012. Elucidating membrane structure and protein behavior using giant plasma membrane vesicles. Nat Protoc 7: 1042–1051.

19. Li, G., J. Kim, Z. Huang, J. R. St Clair, D. A. Brown, and E. London. 2016. Efficient replacement of plasma membrane outer leaflet phospholipids and sphingolipids in cells with exogenous lipids. Proceedings of the National Academy of Sciences of the United States of America 113: 14025–14030.

20. London, E., and G. W. Feigenson. 1978. A convenient and sensitive fluorescence assay for phospholipid vesicles using diphenylhexatriene. Anal Biochem 88: 203–211.

21. Cheng, H. T., Megha, and E. London. 2009. Preparation and properties of asymmetric vesicles that mimic cell membranes: effect upon lipid raft formation and transmembrane helix orientation. The Journal of biological chemistry 284: 6079–6092.

22. Pathak, P., and E. London. 2011. Measurement of lipid nanodomain (raft) formation and size in sphingomyelin/POPC/cholesterol vesicles shows TX-100 and transmembrane helices increase domain size by coalescing preexisting nanodomains but do not induce domain formation. Biophysical journal 101: 2417–2425.

23. St Clair, J. W., and E. London. 2019. Effect of sterol structure on ordered membrane domain (raft) stability in symmetric and asymmetric vesicles. Biochim Biophys Acta Biomembr 1861: 1112–1122.

24. Bakht, O., P. Pathak, and E. London. 2007. Effect of the structure of lipids favoring disordered domain formation on the stability of cholesterol-containing ordered domains (lipid rafts): identification of multiple raft-stabilization mechanisms. Biophysical journal 93: 4307–4318.

25. Levental, I., M. Grzybek, and K. Simons. 2011. Raft domains of variable properties and compositions in plasma membrane vesicles. Proc Natl Acad Sci U S A 108: 11411–11416.

26. Kumar, A., and C. M. Gupta. 1984. Transbilayer distributions of red cell membrane phospholipids in unilamellar vesicles. Biochim Biophys Acta 769: 419–428.

27. Hope, M. J., and P. R. Cullis. 1987. Lipid asymmetry induced by transmembrane pH gradients in large unilamellar vesicles. J Biol Chem 262: 4360–4366.

28. Hope, M. J., T. E. Redelmeier, K. F. Wong, W. Rodrigueza, and P. R. Cullis. 1989. Phospholipid asymmetry in large unilamellar vesicles induced by transmembrane pH gradients. Biochemistry 28: 4181–4187.

29. Lorent, J. H., K. R. Levental, L. Ganesan, G. Rivera-Longsworth, E. Sezgin, M. Doktorova, E. Lyman, and I. Levental. 2020. Plasma membranes are asymmetric in lipid unsaturation, packing and protein shape. Nat Chem Biol 16: 644–652.

30. Keller, H., M. Lorizate, and P. Schwille. 2009. PI(4,5)P2 degradation promotes the formation of cytoskeleton-free model membrane systems. Chemphyschem 10: 2805–2812.

31. Jackson, M. L., and B. J. Litman. 1985. Rhodopsin-egg phosphatidylcholine reconstitution by an octyl glucoside dilution procedure. Biochim Biophys Acta 812: 369–376.

32. Mao, Q., T. Schunk, K. Flukiger, and B. Erni. 1995. Functional reconstitution of the purified mannose phosphotransferase system of Escherichia coli into phospholipid vesicles. J Biol Chem 270: 5258–5265.

33. Dumont, M. E., J. Trewhella, D. M. Engelman, and F. M. Richards. 1985. Stability of transmembrane regions in bacteriorhodopsin studied by progressive proteolysis. J Membr Biol 88: 233–247.

34. Son, M., and E. London. 2013. The dependence of lipid asymmetry upon phosphatidylcholine acyl chain structure. Journal of lipid research 54: 223–231.

35. Suzuki, J., M. Umeda, P. J. Sims, and S. Nagata. 2010. Calcium-dependent phospholipid scrambling by TMEM16F. Nature 468: 834–838.

36. Sharpe, H. J., T. J. Stevens, and S. Munro. 2010. A comprehensive comparison of transmembrane domains reveals organelle-specific properties. Cell 142: 158–169.

37. Brown, D. A., and J. K. Rose. 1992. Sorting of GPI-anchored proteins to glycolipid-enriched membrane subdomains during transport to the apical cell surface. Cell 68: 533–544.

38. Schroeder, R., E. London, and D. Brown. 1994. Interactions between saturated acyl chains confer detergent resistance on lipids and glycosylphosphatidylinositol (GPI)-anchored proteins: GPI-anchored proteins in liposomes and cells show similar behavior. Proceedings of the National Academy of Sciences of the United States of America 91: 12130–12134.

39. Ahmed, S. N., D. A. Brown, and E. London. 1997. On the origin of sphingolipid/cholesterol-rich detergent-insoluble cell membranes: physiological concentrations of cholesterol and sphingolipid induce formation of a detergent-insoluble, liquid-ordered lipid phase in model membranes. Biochemistry 36: 10944–10953.

40. Heerklotz, H. 2002. Triton promotes domain formation in lipid raft mixtures. Biophys J 83: 2693–2701.

41. Dietel, L., L. Kalie, and H. Heerklotz. 2020. Lipid Scrambling Induced by Membrane-Active Substances. Biophys J 119: 767–779.

42. Pantaler, E., D. Kamp, and C. W. Haest. 2000. Acceleration of phospholipid flip-flop in the erythrocyte membrane by detergents differing in polar head group and alkyl chain length. Biochim Biophys Acta 1509: 397–408.

