## Supplemental Figures for "LOSS OF PLASMA MEMBRANE LIPID ASYMMETRY CAN INDUCE ORDERED DOMAIN (RAFT) FORMATION"

Supplemental methods

Thin-Layer Chromatography of PMV Lipids

Lipids in PMV pellets were suspended in 200 μl of PBS and extracted using 2 : 1 (v:v) chloroform : methanol. The lower, lipid-containing chloroform layer was collected, and then 100 μl of water was added, mixed and the aqueous phase was removed. After the lipids were dried with nitrogen, they were dissolved in a small volume of 1 : 1 (v:v) chloroform : methanol and loaded onto a HP-TLC plates (Silica Gel 60), purchased from VWR International (Batavia, IL).) For the TLC analysis two solvent systems were used. The first solvent was 50:38:8:4 (v:v) chloroform : methanol: water : acetic acid. After the solvent front migrated about halfway up the plate, the plate was air-dried for 5 min. Then the plate was re-chromatographed in 1:1 (v:v) hexane : ethyl acetate until the solvent front migrated to near the top of the plate. The lipids were detected by spraying with a 3% (w/v) cupric acetate, 8% (v/v) phosphoric acid solution, then drying for 45 min, and then heating at 180 °C for 2–5 min. For relative quantitation, the TLC plate scanned, PC and SM staining intensities were measured using ImageJ.

Sucrose gradient centrifugation

Sucrose gradient centrifugation was used to confirm if protein and lipid were associated in the OG reconstituted lipid vesicles. 0.4 ml of samples, OG reconstituted PMV (i.e. PMV that were dissolved in OG and vesicles reconstituted by dilution of OG) or reconstituted vesicles from PMV lipids (i.e. vesicles from PMV lipids that were dissolved in OG and then vesicles reconstituted by dilution), which lack protein, were loaded to a 0-20 % (w/w) sucrose gradient centrifuge tube (in which each 0.4ml of 20, 17.5, 15, 12.5, 10, 7.5, 5, 2.5 % sucrose and water were sequentially carefully loaded with highest percentage of sucrose at the bottom). Samples were then centrifuged for 18 h at 37,500 rpm (190,000 x g) at 23°C in a Beckman L8-80 M ultracentrifuge using a SW60 swinging bucket rotor. Fractions of 200 μl were removed sequentially from the top and then diluted to 1 ml with PBS. Tryptophan (Trp) fluorescence in each fraction was measured at λex 280 nm; λem 335 nm. We used the reconstituted sample from PMV lipids (i.e. without protein) as a background. The Trp intensity in each fraction of OG reconstituted PMV was obtained by subtracting the value of fluorescence from the background sample. Then 5 μl of 0.2 mM DPH was added to each fraction, DPH fluorescence was measured to estimate lipid concentration in the vesicles as described in Materials and Methods. Sucrose concentrations were measured using a refractometer, and comparison to standard values.

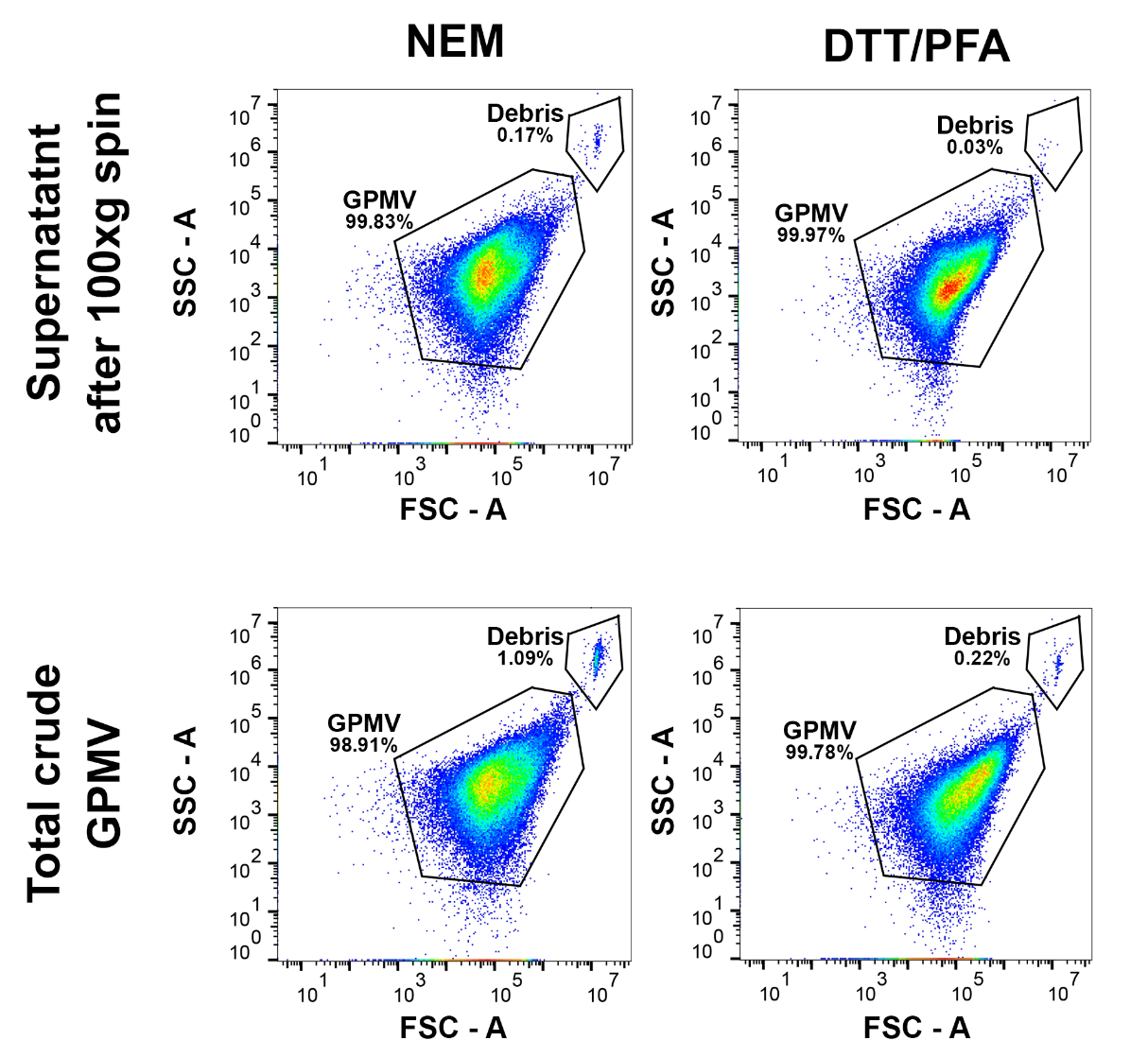

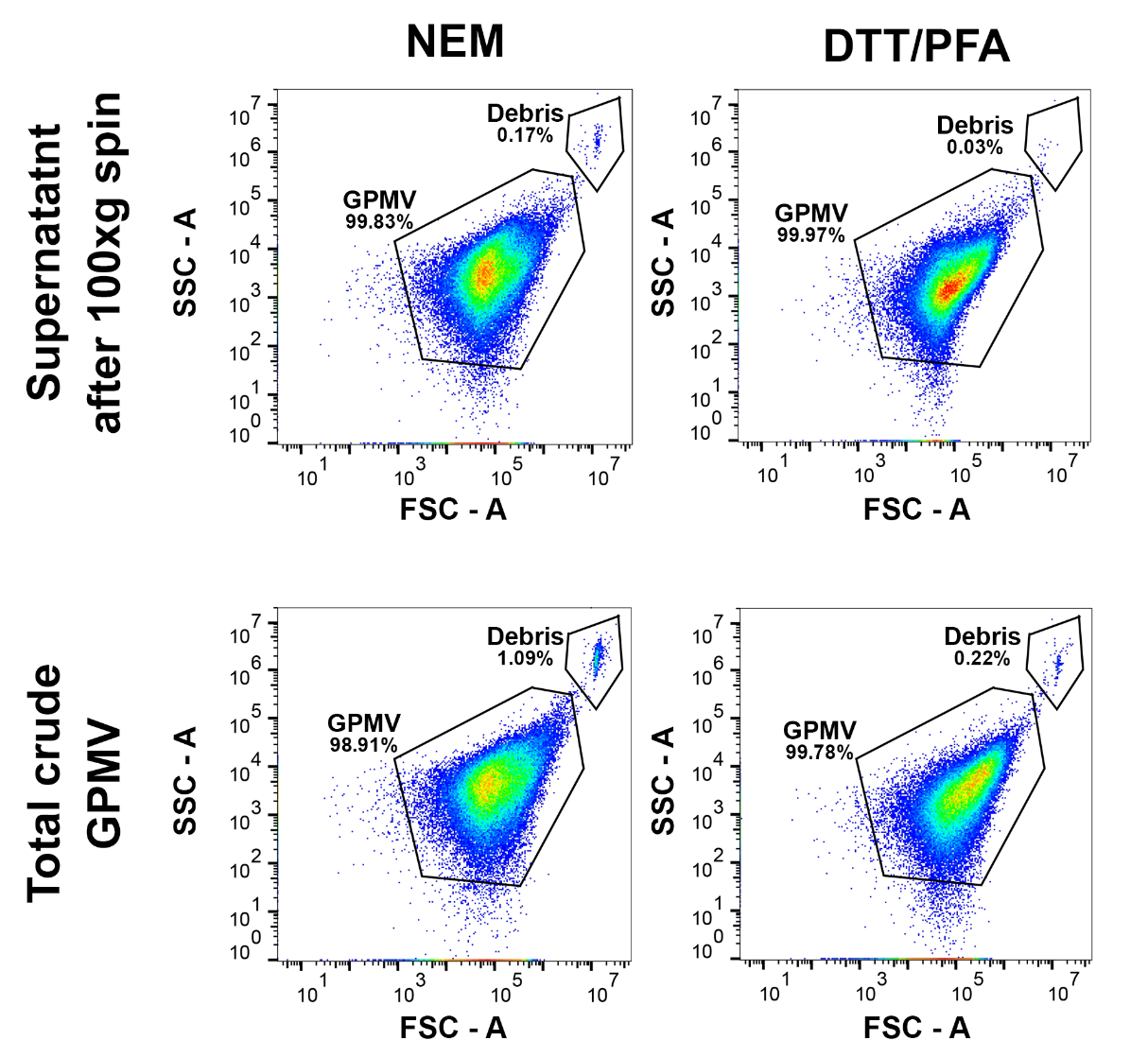

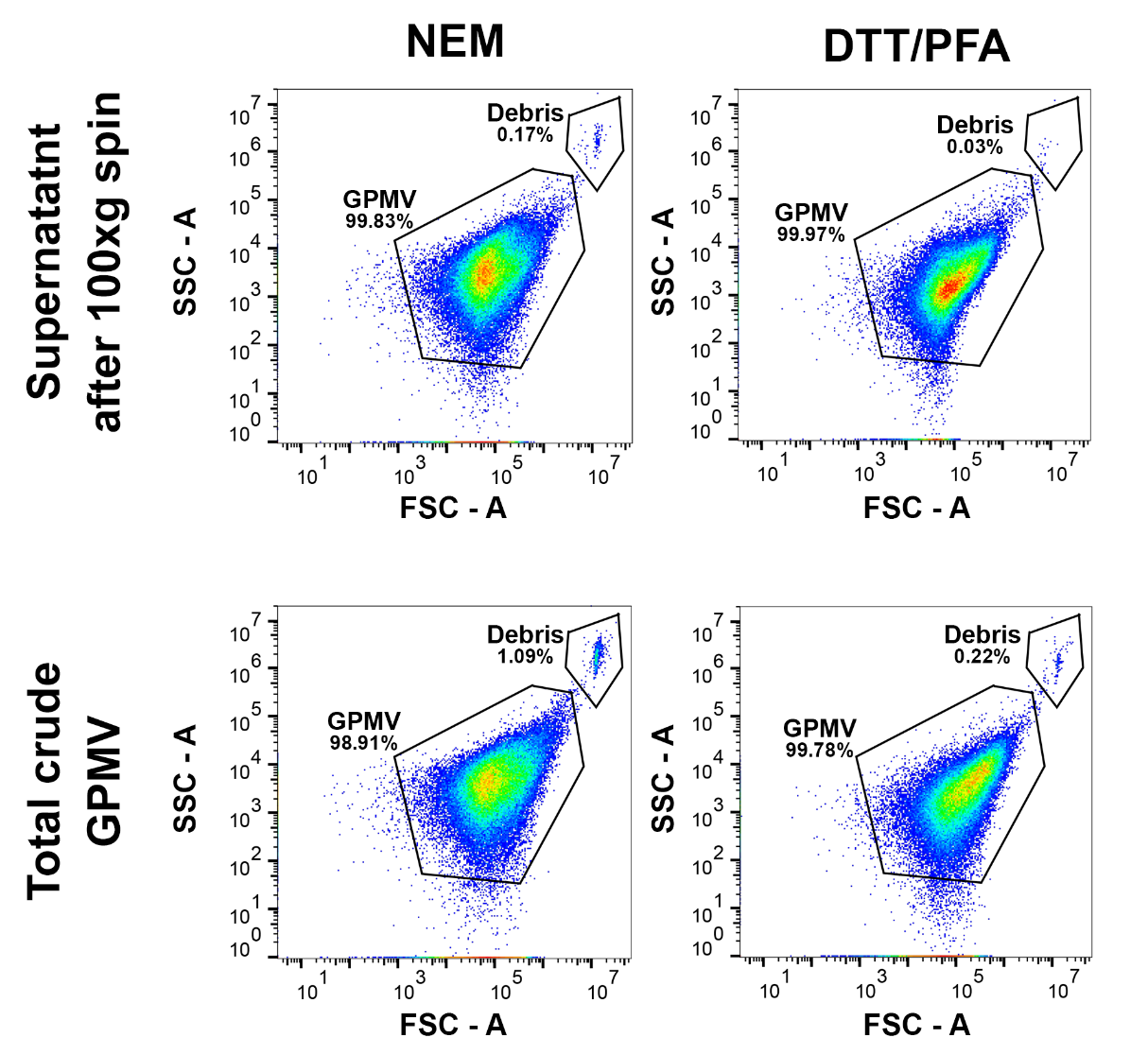

Supplemental Figure 1: Flow cytometry data showing side scatter area (SSC-A) vs. forward scatter area (FSC-A) for GPMVs prepared from RBL-2H3 GPMVs using 2 mM NEM and 5 mM DTT/25 mM PFA. The top panels show the total crude GPMVs obtained from RBL-2H3 cells. The top left and bottom left panels show GPMVs prepared using 2 mM NEM, and the top right and bottom right panels show GPMVs prepared using 5 mM DTT/25 mM PFA. The bottom panels show the 100 x g supernatant containing GPMVs used for annexin binding experiments. The % total gated population in the Debris and GPMVs gates are shown under gate labels.

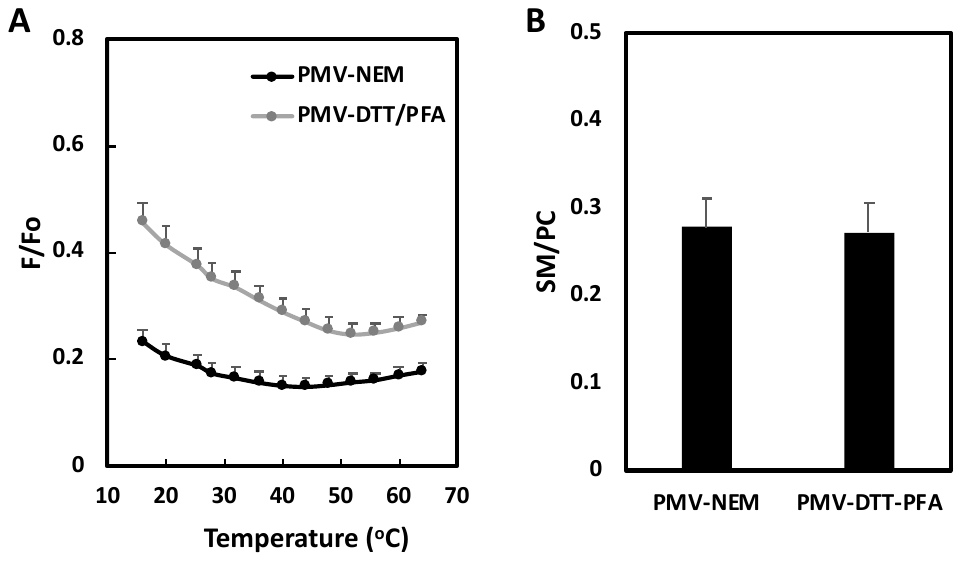

Supplemental Figure 2. (A) Unnormalized F/Fo values for Figure 1A. Figure legend is as in Fig 1A. (B) Analysis of the SM/PC ratio from TLC (as judged by staining intensity). Mean values and standard deviations from three preparations are shown.

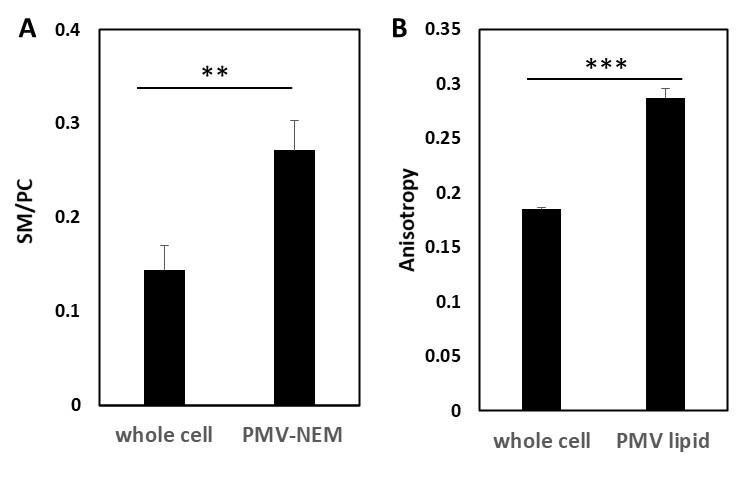

Supplemental Figure 3. Comparison of whole cell and PMV properties. (A) Ratio of SM/PC in whole cell lipids and in PMV preparations as determined by TLC. PMV-NEM = PMV prepared using NEM. (B) Comparison of DPH fluorescence anisotropy in lipid vesicles prepared from whole cell lipids and PMV-NEM lipids at 23 ^o^C. Notice that vesicles formed from whole cell lipids have a lower SM/PC ratio than in vesicles from PMV lipids, and that whole cell lipids form vesicles with less order than vesicles from PMV lipids. Mean values and standard deviations from three preparations are shown. **, *P* < 0.01 , ***, *P* < 0.0005

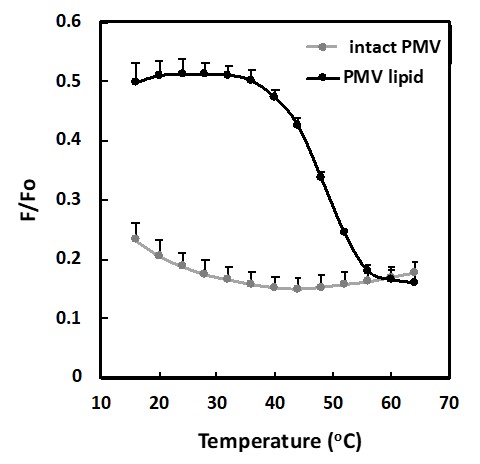

Supplemental Figure 4. Unnormalized F/Fo values for Figure 1B. Figure legend is as in Fig 1B.

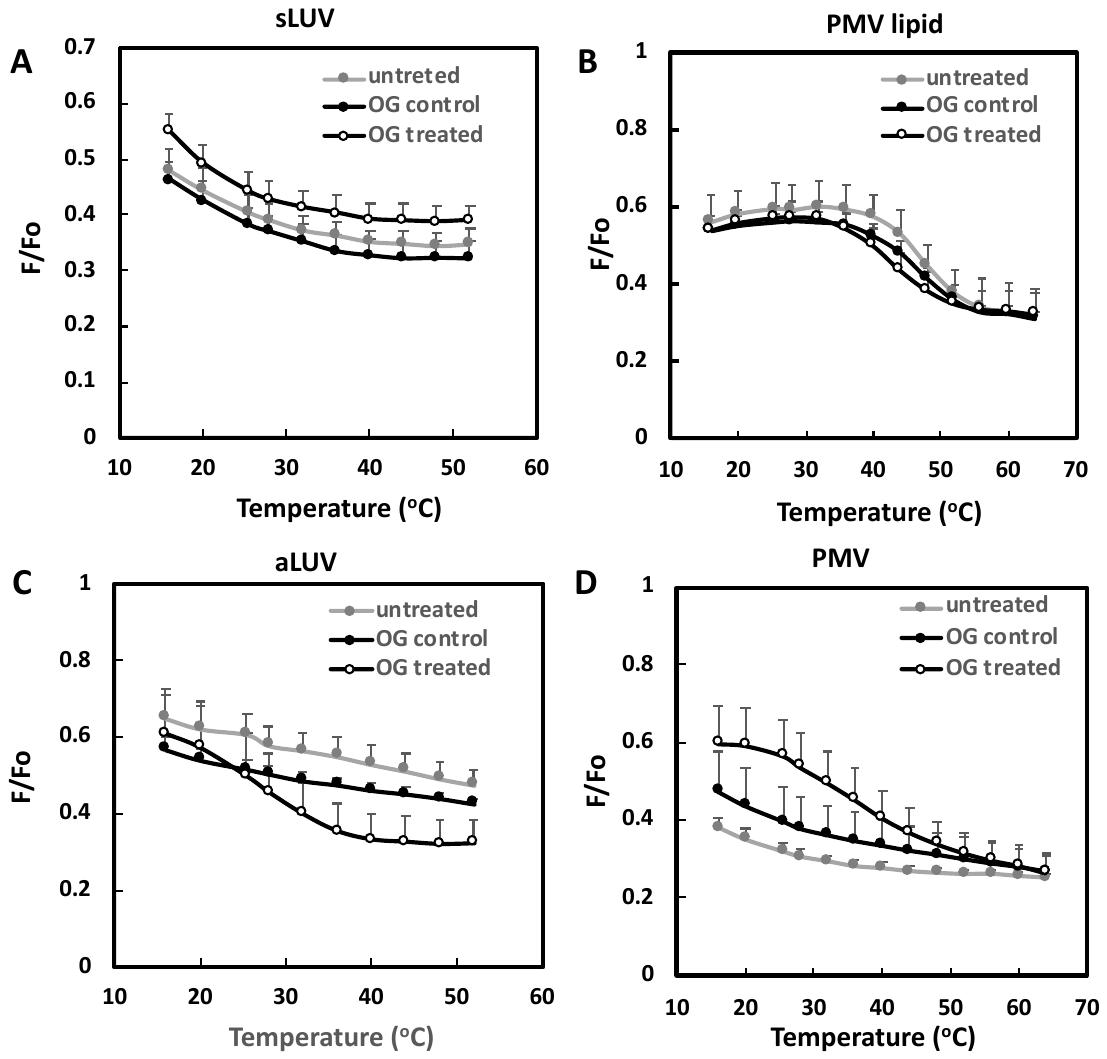

Supplemental Figure 5. Unnormalized F/Fo values for Figure 3. Figure legend as in Figure 3.

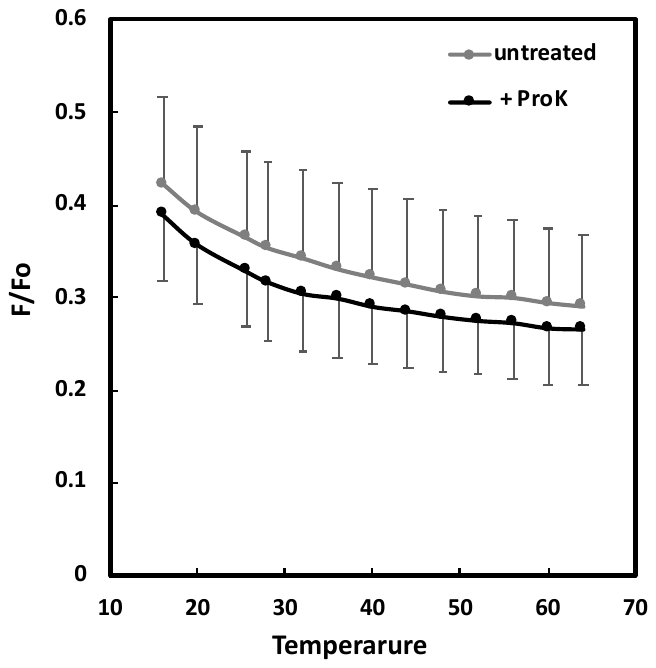

Supplemental Figure 6. Unnormalized F/Fo values for Figure 5A. Figure legend as in Figure 5A.

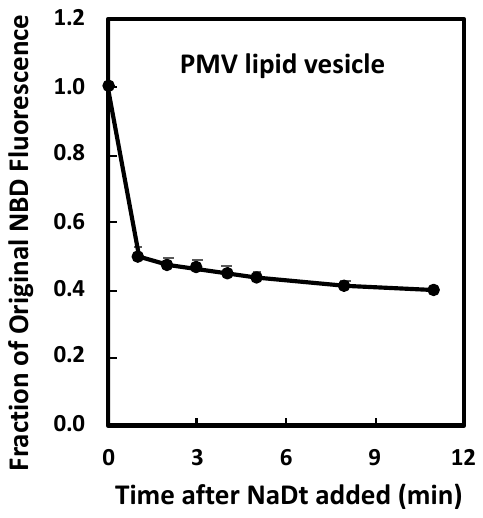

Supplemental Figure 7. Freeze-thaw vesicles from PMV lipids appear to be largely unilamellar. PMV lipid vesicles were prepared with 13.3 μM PMV lipid plus 0.1 mol% of NBD-DPPE, and were dispersed in PBS. At time zero, to a 1 ml sample of lipid vesicles 5 μl of 1 M sodium dithionite (NaDt) dissolved in 1 M Tris pH 10 was added. NBD fluorescence (λex 465 nm, λem 534 nm) was measured vs. time. Extrapolation of fraction of unreacted NBD to time zero indicates close to 50% of the NBD lipid was in the inner leaflet, and so initially inaccessible to NaDt.

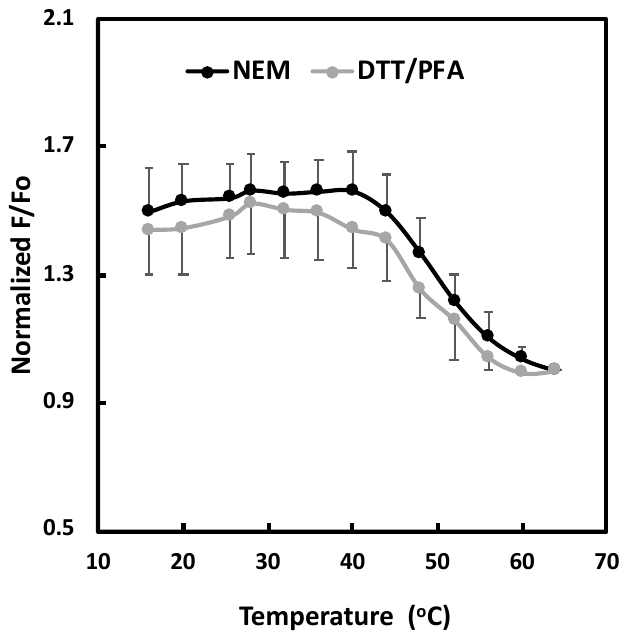

Supplemental Figure 8: Comparison of ordered domain stability for lipid vesicles prepared from NEM and DTT/PFA PMV preparations. Black symbols: Lipids vesicle from PMV lipids prepared using NEM. Gray symbols: Lipid vesicles from PMV prepared using DTT/PFA. Lipid and fluorescence probe concentrations as in Figure 2A.

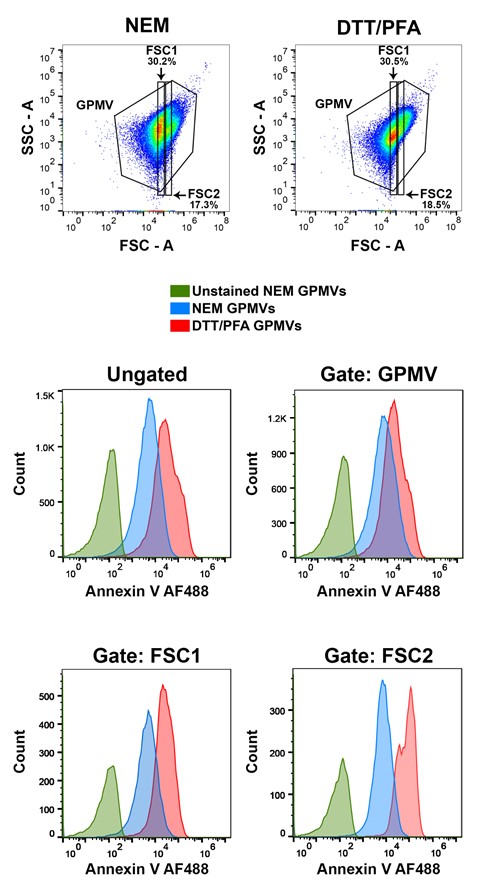

Supplemental Figure 9: Flow cytometry data showing Annexin V AF488 stained GPMVs made using 2 mM NEM and 5 mM DTT/25 mM PFA under different gating conditions. Fluorescence from about 50,000 GPMVs was measured. Top panel shows side scatter area (SSC-A) vs forward scatter area (FSC-A) of GPMVs along with gates: GPMV, FSC1, and FSC2. The % population of FSC1 and FSC2 is shown under gates with respect to the GPMV gate (taken as 100%). The bottom 4 panels show Annexin V-AF488 binding to GPMVs for unstained sample (green), NEM GPMVs (blue), and DTT/PFA GPMVs (red). Unstained DTT/PFA GPMV signal is almost identical to that for unstained NEM GPMV. Middle left panel shows the ungated whole population, middle right panel shows GPMV gate, lower left panel shows FSC1 gate, bottom right panel shows FSC2 gate.

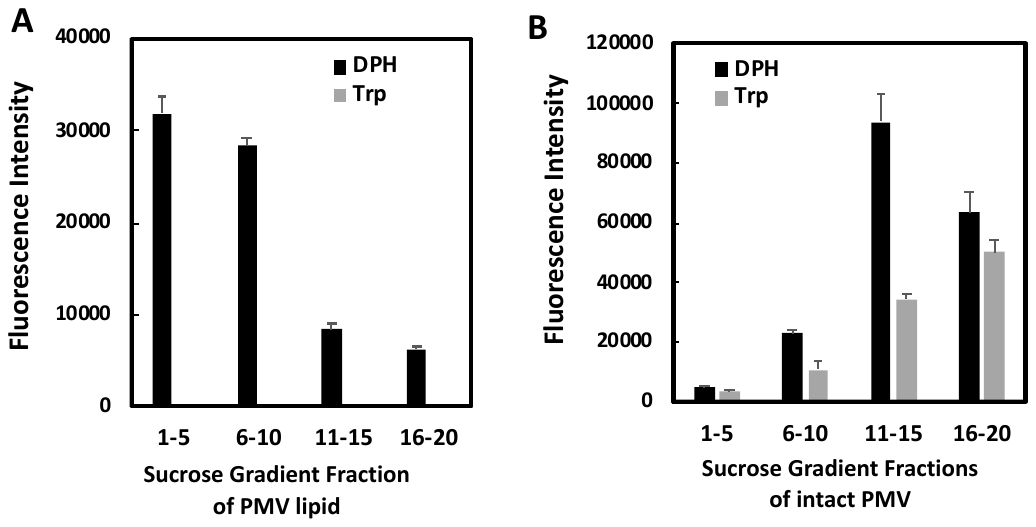

Supplemental Figure 10. Sucrose gradient centrifugation of reconstituted vesicles from PMV lipids or reconstituted PMV. Reconstituted vesicles formed by dissolving vesicles in 45 mM OG and then diluting to 1.5 mM OG. (A) reconstituted vesicles from PMV lipid, and (B) reconstituted vesicles from intact PMV. (black bars) Detection of lipid by measurement of DPH fluorescence. (gray bars) Detection of protein by measurement of Trp fluorescence. (See Materials and Methods for details). Mean relative fluorescence intensities (arbitrary units) and range of duplicates from pooled fractions are shown. Gradient was roughly linear from 2-20 w/w % sucrose. Approximate sucrose concentrations from index of refraction for pooled fractions 1-5, 6-10, 11-15 and 16-20 were 3, 7.0, 12.5, and 17 w/w % sucrose, respectively.

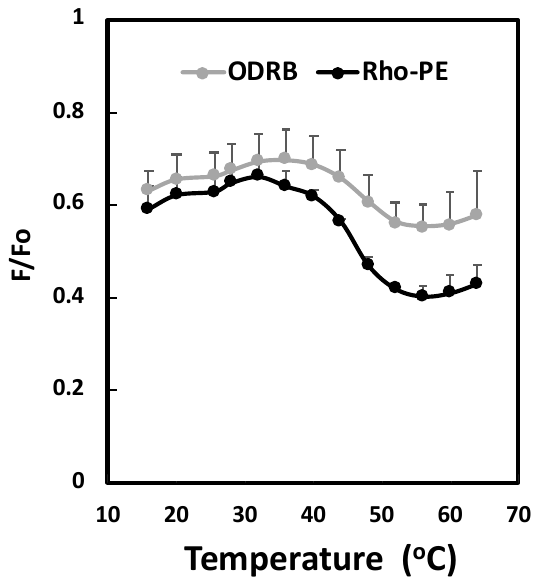

Supplemental Figure 11. Comparison of FRET for lipid vesicles prepared from PMV lipid extracts when FRET acceptor is ODRB or Rhodamine-DOPE (Rho-PE). Lipid vesicles dispersed in PBS. Lipid concentration was 20 μM. ODRB concentration was 0.4 μM, rhodamine-DOPE concentration was 4 mol% of lipid (0.8 μM). DPH concentration was 0.03 μM .
